# How do brain regions specialised for concrete and abstract concepts align with functional brain networks? A neuroimaging meta-analysis

**DOI:** 10.1101/2024.07.11.603079

**Authors:** Paul Hoffman, Matthew Bair

## Abstract

Identifying the brain regions that process concrete and abstract concepts is key to understanding the neural architecture of thought, memory and language. We review current theories of concreteness effects and test their neural predictions in a meta-analysis of 72 neuroimaging studies (1400 participants). Our analysis includes more than twice as many studies as previous meta-analyses, allowing for a more sensitive mapping of these effects across the brain. We also conducted a quantitative assessment of the degree to which concreteness effects aligned with a range of large-scale functional brain networks. Our results suggest that concrete and abstract concepts vary both in the information-processing modalities they engage and in the demands they place on cognitive control processes. Abstract concepts preferentially activated networks for social cognition (particularly for sentences), language and semantic control (particularly when presented as single words). Concrete concepts preferentially activated action processing regions when presented in sentences, though we found no evidence that they activated visual networks. Specialisation for both concept types was present in different parts of the default mode network (DMN), with effects dissociating along a social-spatial axis. Concrete concepts generated greater activation in a medial temporal DMN component, implicated in constructing mental models of spatial contexts and scenes. In contrast, abstract concepts showed greater activation in frontotemporal DMN regions involved in social and language processing. These results align with prior claims that generating models of situations and events is a core DMN function and indicate specialisation within DMN for different aspects of these models.

## Introduction

Human language and experience encompasses a vast array of concepts along a spectrum of concreteness. At one end of the spectrum lie highly concrete words that label objects and acts we directly perceive in our environment (e.g., *a parrot, to jump*). At the other, we find highly abstract words whose meanings are intangible and divorced from direct sensorimotor experience (e.g., *a theory, to believe*).^1^ Accounting for this conceptual diversity is a central challenge for theories of language comprehension and semantic cognition (Reilly et al., 2025). A range of distinct theories have been developed to explain processing differences between concrete and abstract concepts, motivated by a large body of behavioural and neuroscientific data. In this article, we review these theories and identify their predictions about how concrete and abstract concepts differentially activate the cortex. We then test these predictions in meta-analysis of 72 neuroimaging studies (1400 participants) that contrasted activation to concrete and abstract language. Our study represents a significant advance on previous meta-analyses of these effects in three ways. First, it includes twice as many studies as previous meta-analyses (Binder et al., 2009; Bucur & Papagno, 2021; J. Wang et al., 2010), allowing for a more sensitive mapping of these effects across the brain. Second, we tested for the first time how neural concreteness effects vary systemically along theoretically-important stimulus and task dimensions. Third, we conducted a quantitative assessment of the degree to which concreteness effects aligned with a range of large-scale functional brain networks. By cleaving the neural response to meaning along the fundamental dimension of concreteness, we gain new insights into the ways in which the brain constructs representations of the different forms of experience conveyed through language.

Many of the major accounts of concreteness effects stem from an embodied (or grounded) approach to semantics (Barsalou, 1999; Gallese & Lakoff, 2005; Glenberg & Robertson, 2000; Pulvermüller, 2013). Developed in opposition to classical symbolic views of language (Johnson-Laird, 1988; Newell, 1980), the embodied view holds that conceptual representations are closely tied to the perceptual experiences they reference and that understanding a concept involves reactivating those experiences in some way. The embodied semantics umbrella shelters a broad range of specific theories, from strong accounts that emphasise the primacy of embodied reactivation of sensorimotor cortices (Gallese & Lakoff, 2005; Glenberg & Robertson, 2000) to those that propose a central role for multimodal hub regions that bind and integrate the various types of experience associated with each concept (Binder, 2016; Binder & Desai, 2011; Lambon Ralph et al., 2017). These theories share the general assumption that understanding words frequently involves activating neural networks that process the experiences the words refer to.

Embodied semantics accounts have traditionally focused on the involvement of sensorimotor modalities in semantic representation. An early forerunner of these, Dual-Coding theory, proposed that concrete words are represented in a visual semantic system in addition to a verbal system shared by abstract words (Paivio, 1986). More recently, evidence has accumulated for activation of a range of sensorimotor networks during concrete word processing. Motor and motion-processing regions are activated when we process language that describes actions (Giacobbe et al., 2022; Hauk et al., 2004; Saygin et al., 2010; Watson et al., 2013) and parts of the ventral visual stream are activated when people make judgements about the shapes and colours of words’ referents (Fernandino et al., 2016; Martin et al., 1995; Thompson-Schill et al., 1999; van Dam et al., 2012). This embodied perspective makes a straightforward prediction about concreteness effects in the brain. Since concrete words are more strongly associated with sensorimotor experiences, they should activate sensorimotor cortices to a greater extent. As vision is the dominant modality influencing a word’s concreteness (Connell & Lynott, 2012), we would expect strong concrete > abstract (C>A) activation effects in the visual system. Action processing cortex should show similar effects, particularly for sentences that contain action verbs.

The representation of abstract concepts has frequently been presented as a challenge to the embodied semantics approach, because they do not directly reference sensorimotor experiences. This challenge has been addressed in a number of ways. Conceptual Metaphor Theory (Gibbs, 1994; Lakoff & Johnson, 1999) asserts that abstract concepts activate sensorimotor representations indirectly through metaphor and analogy (e.g., the association of power with vertical height) (Pecher, 2018). Few studies have investigated whether abstract concepts activate sensorimotor cortices indirectly via metaphoric association (Desai, 2022; Joue et al., 2020) but it seems likely that such indirect activation would be weaker than that elicited by concrete concepts. Thus, this view is also compatible with C>A effects in sensorimotor cortex.

Other researchers have considered other forms of experience that abstract words draw upon. The Affective Embodiment view (Kousta et al., 2011; Lenci et al., 2018; Newcombe et al., 2012; Vigliocco et al., 2014) holds that abstract concepts are more closely associated with emotional content than concrete words (but see (Winter, 2023)) and should therefore preferentially engage brain regions involved in experiencing emotions. This perspective is part of a wider move to ground semantics in activation of internal bodily states and experiences as well as in the external world (Barsalou, 2008; Connell et al., 2018; Martin, 2016). A related view emphasises the importance of social experiences in the acquisition and understanding of abstract concepts (Borghi et al., 2019; Pexman et al., 2022; Troche et al., 2014; Wiemer-Hastings & Xu, 2005). Many abstract words refer to social interactions at an interpersonal or societal level (e.g., loan, democracy) and might therefore be supported by simulations of our own or others’ mental states. These related views predict abstract > concrete (A>C) activation effects in brain regions that process emotion and regions involved in social cognition and theory of mind (Andrews-Hanna et al., 2014; Yeshurun et al., 2021).

In addition to direct experience, we gain knowledge about concepts through their expression in language. The successes of large computational language models have demonstrated that sophisticated semantic information can be derived solely from the statistical patterns of natural language use (J. Devlin et al., 2019; Mikolov et al., 2013; Pereira et al., 2016). In line with this position, a number of hybrid theories suggest that semantic processing is supported by language-based representations as well as by simulations in modal brain systems (Barsalou et al., 2008; Binder & Desai, 2011; Connell, 2019; Dove et al., 2022; Louwerse, 2011; Vigliocco et al., 2009). Verbal experience is more critical to the acquisition of abstract words and is thought to play a more central role in determining their meanings (Borghi et al., 2019; Vigliocco et al., 2009; Villani et al., 2019). These hybrid views therefore predict A>C effects in temporal lobe regions involved in language comprehension. Some researchers see linguistic representations of meaning as most suited to supporting superficial processing (Barsalou et al., 2008; Connell, 2019). A>C effects in language cortex might therefore be most prominent for simple tasks, while deeper conceptual processing instead requires more complex representations grounded in other brain systems.

Thus far, we have focused on the types of experience that contribute to our understanding of concrete and abstract words. Other accounts have investigated the role of context in shaping our understanding of each concept type. It has long been known that when people are presented with isolated abstract words, they find it hard to generate contexts in which the word could be used (Schwanenflugel et al., 1988; Schwanenflugel & Shoben, 1983). This is probably because abstract words are used in a wider variety of contexts than concrete words, suggesting that their meanings are highly context-dependent (Davis et al., 2020; Hoffman et al., 2013). Abstract word meanings also vary greatly between individuals (X. Wang & Bi, 2021) and are difficult to process in the absence of context (Bechtold et al., 2021; Hoffman et al., 2010; Schwanenflugel et al., 1988; Schwanenflugel & Shoben, 1983). A related line of work proposes that concreteness is correlated with conceptual specificity: abstract words tend to have less specific referents than concrete words (Villani et al., 2024). In other words, the semantics of abstract words are frequently vague, volatile and contextually-varying. Because of this inherent uncertainty, we and others have proposed understanding abstract words relies more on executive control processes that regulate how semantic knowledge is activated (Bechtold et al., 2021; Cousins et al., 2016; Davis et al., 2020; Hoffman, 2016; Hoffman et al., 2010, 2015). These accounts predict A>C activation in brain regions that support controlled semantic processing (Jackson, 2021) and predict that these effects should be largest (a) when no context is available to support processing (i.e., for single words rather than sentences) and (b) when participants are required to make a specific semantic judgement about the stimulus.

Finally, a related account proposes that abstract concepts vary greatly in the real-world situations or events that they describe, as well as in their verbal contexts (Barsalou et al., 2018; Davis et al., 2020). For example, an abstract concept like *justice* could sometimes bring to mind a courtroom situation and at other times a prison (Davis et al., 2020). It has also been suggested that we rely heavily on situational features – i.e., the spatiotemporal context of an experience, the entities present and their interactions – to bring meaning to abstract concepts (Barsalou et al., 2018; Barsalou & Wiemer-Hastings, 2005; Binder, 2016; Davis et al., 2020; McRae et al., 2018). This proposal is related to the claim that social interactions are highly relevant to abstract words (Borghi et al., 2019; Pexman et al., 2022; Troche et al., 2014) and that abstract concepts are structured by their thematic relationships with one another (Crutch & Warrington, 2005). It is hard to derive clear predictions about neural concreteness effects from this situational account. If processing abstract words involves simulating the situations they could refer to, we might expect A>C activation in brain regions that represent situation models (Ranganath & Ritchey, 2012; Yeshurun et al., 2021). However, given that abstract concepts are weakly associated with a wide range of possible situations, such effects might only occur when abstract words are placed in a specific constraining context (e.g., a sentence). During processing of single words, we might expect concrete words to activate situational cortex more, since these words bring specific situations to mind more readily (Binder, 2016; Schwanenflugel et al., 1988).

As this brief review has shown, identifying neural processing differences between concrete and abstract concepts is critical to unlocking a deeper understanding of how we bring meaning to experience and language. The various theoretical claims and their neural predictions are summarised in Table 1. To test these, we performed quantitative co-ordinate-based meta-analyses of 72 neuroimaging studies that manipulated the concreteness of language stimuli. We formally assessed how concreteness effects intersect with the brain regions that support 7 key processing domains (see Figure 2). By examining the differential contributions of these systems to concrete and abstract processing, we were able to assess the degree of support for each theoretical position. These theories also make specific predictions about concreteness effects depend on the stimuli and tasks used to elicit them. Thus, we systematically investigated how the expression of C>A and A>C effects is influenced by two theoretically-relevant factors: (a) the distinction between single-word and sentence stimuli and (b) the depth of semantic processing participants were required to engage in.

**Table 1:** Summary of predictions about concreteness effects and degree to which they were supported in the meta-analysis.

| **Claim** | **C>A predictions** | **Supported?** | **A>C predictions** | **Supported?** |
| --- | --- | --- | --- | --- |
| Understanding concrete language is more reliant on activating its sensorimotor correlates | Action network<br>Visual network | For sentences<br>No |  |  |
| Understanding abstract language is more reliant on activating its social and emotional associations |  |  | Social network<br>Emotional network | Yes, especially sentences<br>No |
| Understanding abstract language is more reliant on activating linguistic knowledge |  |  | Language network | Yes, especially semantic tasks |
| The meaning of abstract language is more variable than that of concrete language |  |  | Semantic Control | Yes, especially single words |
| Situation models can be more engaged when processing concrete or abstract language, depending on the level of contextual support | Situation network | Yes, especially semantic tasks | Situation network? | For sentences |

We also assessed where concreteness effects occur within the default-mode network (DMN), which appears to play an important role in generating internal mental models of experiences. DMN is often deactivated for tasks that require sole focus on the external environment, leading to its traditional characterisation as a task-negative network (Buckner et al., 2008; Raichle et al., 2001). However, tasks that require use of internally-generated representations, including spontaneous thoughts, episodic memories and semantic knowledge, positively engage this network (Andrews-Hanna, 2012; Binder et al., 2009; Ranganath & Ritchey, 2012). These findings, along with the observation that DMN regions are convergence points for multiple modal processing streams (Binder & Desai, 2011; Margulies et al., 2016), have led to proposals that DMN integrates incoming information with past experience to mentally simulate complex multimodal events and situations (Baldassano et al., 2018; Fernandino & Binder, 2024; Heidlmayr et al., 2020; Morales et al., 2022; Ranganath & Ritchey, 2012; Yeshurun et al., 2021). DMN regions are therefore prime candidates for supporting the different mental simulations required to understand concrete and abstract concepts. The DMN can be fractionated into three sub-systems with distinct functional profiles (Andrews-Hanna et al., 2010, 2014; Lee et al., 2021; Shao et al., 2024; Yeo et al., 2011), shown in Figure 2. First, a medial temporal sub-network (MT-DMN) encompasses the hippocampus and adjacent medial temporal cortex, as well as parts of retrospenial, posterior inferior parietal and ventromedial prefrontal cortices. This subnetwork is particularly involved in episodic memory and the mental representation of scenes and spatial contexts. Second, a frontotemporal system (FT-DMN; sometimes termed dorsal medial DMN) consists of lateral temporal, dorsomedial prefrontal and anterior inferior parietal cortices, as well as lateral prefrontal regions that frequently co-activate with them. This subnetwork has been implicated in semantic and social cognition. Finally, a core system (core-DMN) is spatially interposed between the other two systems and shows some functional characteristics of both, but with particular engagement for self-relevant processing. It has recently been suggested that these systems might differentially support concrete and abstract aspects of thought and imagination (Andrews-Hanna & Grilli, 2021). Specifically, the complex semantic representations of abstract words and their reliance on social cognition might lead these regions to preferentially activate FT-DMN regions, while the grounding of concrete concepts in external spatial contexts might associate these with the MT-DMN system. Our meta-analysis assesses this proposal within the domain of language processing for the first time.

## Method

### Study selection

The process for identifying studies is summarised in Figure 1. We searched for peer-reviewed studies published between January 1990 and August 2022, which contrasted processing of concrete and abstract language. An initial search was conducted using the Scopus database for articles containing the following terms in their title, abstract or keywords: ["concrete*" OR "abstract*" OR "imageab*"] AND ["fMRI" OR "PET"] AND ["word*" OR "language" OR "semantic*" OR "sentence*" OR "memory"]. This yielded 903 studies, which were screened for inclusion.

**Figure 1:**
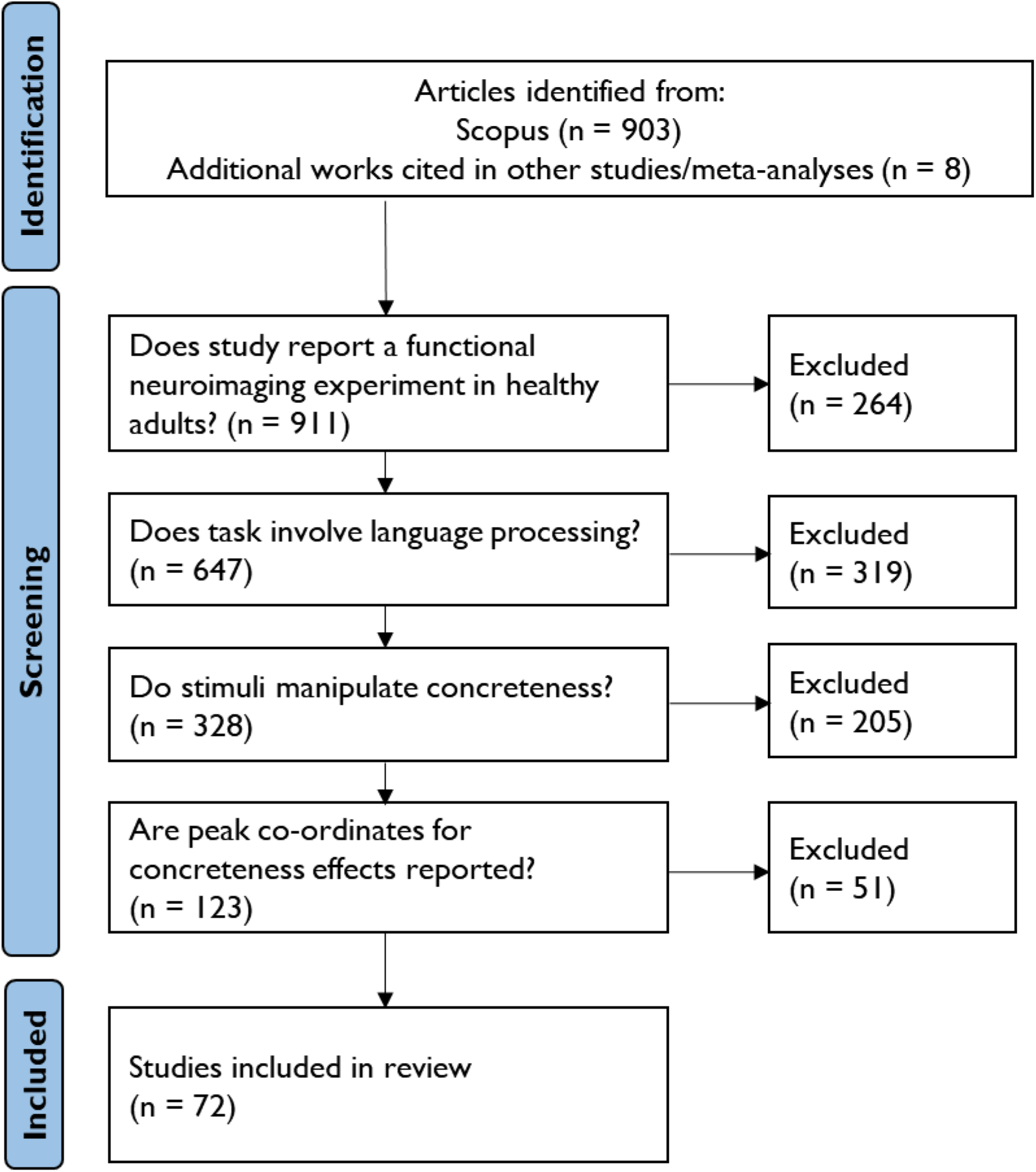
Process used to identify studies.

The inclusion criteria were:

1. The article reported the results of a functional neuroimaging experiment in healthy adult participants.
2. Participants were required to process meaningful language stimuli in their native language, which could be presented in written form or as audio or video recordings. We placed no restriction on the type of processing required.
3. The language stimuli varied in concreteness, either in a factorial design (concrete vs. abstract) or through a parametric manipulation of concreteness values. We included all studies where the researchers referred to their conditions as concrete and/or abstract. In most cases, researchers reported that their conditions differed in concreteness or imageability ratings collected for the study or obtained from a published database, such as (Brysbaert et al., 2014).
4. The article reported activation peaks for a univariate contrast between concrete and abstract conditions (or effect of parametric variation in concreteness) in a standard stereotactic space (MNI or Talairach).

A total of 64 studies were identified through this process. We added to this 8 further studies that met the inclusion criteria but were not returned in the Scopus search. These were either cited by studies that passed the initial screening process and were present in previous neuroimaging meta-analyses of concreteness effects (Binder et al., 2009; Bucur & Papagno, 2021; J. Wang et al., 2010). Thus, a total of 72 studies were included in analyses (see Supplementary Materials for a full list). The vast majority of these studies took a factorial approach, labelling one set of stimuli concrete and the other abstract, rather than treating concreteness as a continuous variable. Thus, our meta-analysis follows the same approach.

Studies varied in whether they defined concrete and abstract categories on the basis of concreteness ratings (how much does the word refer to something that can be experienced through the senses?) or imageability ratings (how easily does the word elicit a mental image?). These are very strongly correlated with one another (*r >* 0.8), though both are biased towards visual experience (Connell & Lynott, 2012). An alternative approach is to ask participants to rate association with sensory experience in various modalities separately and take the maximum rating as a word’s perceptual strength (Connell & Lynott, 2012). However, no neuroimaging studies have taken this approach to classifying words as concrete or abstract. Many studies attempted to match their stimuli on other properties that might influence brain activity (e.g., word/sentence length, word frequency). However, there was considerable heterogeneity in which properties researchers chose to equate, reflecting the diverse range of research questions under investigation. This heterogeneity means that the meta-analysis was able to identify concreteness effects that hold across a range of different study designs that control for different confounding factors.

To investigate how concreteness effects were influenced by relevant study characteristics, we classified studies in two dimensions.

1. <u>Stimulus.</u> *Word* studies used single-word stimuli (including those that presented pairs or triads of single words for semantic decisions). *Sentence* studies used sentence-level stimuli (including three studies that used noun-verb phrases).
2. <u>Task.</u> *Semantic* studies used tasks that explicitly required participants to make a judgement based on the meaning of the stimuli (e.g., semantic relatedness judgements, pleasantness judgements, sensibility judgements). *Non-semantic* studies did not require explicit meaning-based decisions (e.g., passive reading/listening, lexical decision, memory tasks).

For both of these dimensions, where a study could not be classified unambiguously it was assigned to neither category.

### ALE analyses

Activation Likelihood Estimation (ALE) analyses were carried out using GingerALE 3.0.2 (Eickhoff et al., 2012). ALE is the most commonly-used method for co-ordinate-based meta-analysis of neuroimaging data. The software takes peaks from neuroimaging contrasts of interest across a set of studies, models the spatial distribution of these peaks and computes activation likelihood maps for the whole brain. These maps are then subjected to voxel-wise statistical tests to identify regions that consistently show effects across studies.

We first used the ALE method to identify regions showing greater activation for concrete relative to abstract words (C>A) and the reverse (A>C) in the full set of 72 studies. To investigate the factors influencing these effects, we then conducted subtraction analyses that contrasted studies on the two dimensions. For example, the Stimulus C>A subtraction analyses identified regions that were more likely to show C>A effects for Word studies than in Sentence studies and vice-versa. There were four subtraction analyses in total (C>A and A>C for Stimulus and Task contrasts).

We used Turkeltaub et al.’s (2012) non-additive ALE algorithm, which limits the influence of individual studies reporting multiple peaks close to one another. Peaks reported in Talairach space were converted to MNI space using the tal2icbm_spm transform (Lancaster et al., 2007). The main C>A and A>C results were thresholded using the recommended permutation-based method for cluster-level inference (Eickhoff et al., 2012). A family-wise error cluster-corrected threshold of *p* < 0.05 was used (with a cluster-forming threshold of p < 0.005 and 10,000 permutations; Hoffman & Morcom, 2018). Cluster-level inference is not available for subtraction analyses so for these we used a voxelwise threshold of *p* < 0.05 combined with a minimum cluster extent of 1600mm^3^ and 100,000 permutations.

### Comparison with functional brain networks

We used three methods to systematically interpret the functional correlates of regions exhibiting concreteness effects. First, we submitted unthresholded ALE maps to the Neurosynth website (Neurosynth.org) to obtain correlations with its meta-analytic probability maps. Neurosynth is an automated meta-analysis tool that identifies terms commonly used in a large corpus of neuroimaging papers (14,371 studies for the analyses reported here) and relates these to reported peak activation co-ordinates in the same papers. A Neurosynth map for a given term indicates the likelihood that activation in each voxel is preferentially reported by studies that use the term. For example, in the map for “language”, each voxel’s value indicates the likelihood that activation is reported there in papers that discuss language, relative to studies that don’t. This large-scale unsupervised approach to meta-analysis has been validated by showing that it can successfully replicate known functional networks identified in manual meta-analyses (Rubin et al., 2017; Yarkoni et al., 2011). It is important to note, however, that the automated nature of Neurosynth makes it unsuitable for investigating concreteness effects. Papers associated with the term “concreteness” usually report both C>A and A>C peaks and Neurosynth’s algorithms do not discriminate between these.

By correlating the Neurosynth term maps with our unthresholded ALE maps, we were able to determine which terms are most consistently associated with regions that show concreteness effects. For the main analysis, we subtracted A>C ALE values from C>A ALE values and correlated the result with Neurosynth maps. Here, positively-correlated terms were more associated with regions showing C>A effects and negatively-correlated terms with A>C effects. To visualise the results, terms relating to anatomical structures were removed and the 30 most strongly correlated terms in each direction were extracted. These were plotted as word clouds in which the size of each term is proportional to the strength of its correlation with concreteness effects. A similar process was followed to visualise associations with the effects of each study dimension.

Second, we compared the spatial distribution of concreteness effects with a series of functional brain networks. We selected networks that support the various processes that concrete and abstract words have been hypothesised to differentially rely on, again mostly based on Neurosynth meta-analytic maps. These were:

1. Action. Regions associated with the “action” term.
2. Visual. Regions associated with the “visual” term.
3. Emotions. Regions associated with the “emotional” term.
4. Social. Regions associated with the “social” term.
5. Language. Regions associated with the “language comprehension” term.
6. Semantic control. No term similar to semantic control was present in the Neurosynth database so we instead used the network identified in a recent meta-analysis of studies manipulating controlled processing demands in semantic tasks (Jackson, 2021).
7. Situations. “Situations” was not present in the Neurosynth database as an individual term. Instead, we searched Neurosynth’s topic-based maps (Rubin et al., 2017) for a topic that related to processing of situations and events. We selected a topic from the v5-topics-200 set for which the highest-loading terms were: events, future, personal, past. We reasoned that these terms index tasks in which situation models are hypothesised to play an important role (i.e., thinking about events, remembering the past, contextualising oneself and imagining the future).

Neurosynth maps (z-values for test of the association between voxel activation and term use, thresholded at FDR-corrected *p* < 0.01) were downloaded from neurosynth.org (v5 dataset), smoothed at 4mm FWHM and binarized at a threshold of *z*>3. The map of significantly-activated regions for semantic control was downloaded from https://github.com/JacksonBecky/SemanticControlMetaA and binarised. Finally, we compared the spatial distribution of concreteness effects across sub-systems of the default mode network (DMN). Following other researchers (Andrews-Hanna et al., 2014; Poerio et al., 2017; Shao et al., 2024), we used Yeo et al.’s (2011) resting-state parcellation of the brain into 17 networks to define frontotemporal, core and medial temporal DMN sub-networks.

To determine the degree to which concreteness effects aligned with the functional and DMN networks, we computed the Jaccard similarity between each network and the regions showing significant effects in each ALE analysis. Jaccard similarity quantifies the degree to which two analyses implicate the same brain regions, with 1 indicating that significant regions in both analyses are identical and 0 indicating no regions in common (Maitra, 2010). Of course, some overlap between networks can occur by chance, particularly when the networks being considered cover large areas of the cortex. Thus, we used a permutation approach to determine when the overlap between ALE results and networks was greater than expected by chance. Using the Neuromaps toolbox (Markello et al., 2022), each network and ALE result was projected to the fsaverage5 cortical surface. For each ALE analysis, we then generated 10,000 null versions of the data by inflating the surface to a sphere and rotating it by a random amount (the spin test; Alexander-Bloch et al., 2018). This method preserves the spatial structure within the effect map but randomises its position on the cortical surface. We computed the Jaccard similarity of each null ALE result to each network, resulting in distributions of expected similarities under the null hypothesis. The position of the true Jaccard similarity value in its corresponding null distribution was used to generate a p-value for the overlap. P-values for each analysis were corrected for multiple comparisons using the false discovery rate (FDR) approach.

### Data availability

The raw data for these analyses, including all peak co-ordinates and the studies and contrasts they were obtained from, are available on the OSF website: https://osf.io/78ra9/?view_only=1104705db40e4bfaa0744177d6563213. Unthresholded ALE maps for all analyses are available at the same address, as is the code used to perform the network comparisons.

## Results

### Literature Search

We identified 72 studies that reported peak activation co-ordinates for contrasts of concrete vs. abstract language stimuli. Studies were published between 1995 and 2022 and together comprised 10 languages, 1400 participants, 459 C>A peaks and 435 A>C peaks. The number of included studies is considerably higher than older meta-analyses on this topic (Binder et al., 2009; J. Wang et al., 2010). It is also higher than the 32 studies analysed more recently by Bucur & Papagno (2021), principally because these authors only included studies that used single-word stimuli (whereas we also included phrases and sentence stimuli). This increase in scale provides a number of advantages. First, our analysis includes results from more recent studies which tend to use larger sample sizes than the older (pre-2010) literature and thus are sensitive to smaller effects. Second, by sampling a larger number of studies, we were able to perform comparisons of subsets of studies with different characteristics. Third, the inclusion of a wide variety of studies means there is greater heterogeneity in the study designs and procedures that generated the data. As ALE analyses treat study as a random effect, this heterogeneity means that the analysis identifies effects that generalise over a wide range of conditions.

As reported in Supplementary Table 1, studies used a range of tasks to contrast concrete and abstract words, including passive reading/listening, lexical decision, semantic decisions of various types and memory encoding and retrieval tasks. A range of stimuli were also employed, with some studies presenting individual words and others presenting phrases or complete sentences. To investigate how these study characteristics influenced the expression of concreteness effects, we classified studies along two theoretically-relevant dimensions: *Stimuli* (23 Sentence studies; 48 Word studies; 1 unclassifiable) and *Task* (30 Semantic; 40 Non-semantic; 2 unclassifiable). There was no statistical association between these two dimensions (*χ*^2^(1) = 0.18, *p* = 0.67).

### Elucidation of relevant cortical networks

To aid interpretation of our results, we first defined 7 relevant functional networks (obtained from existing meta-analytic resources; see Method for details) and 3 sub-divisions of the DMN, obtained from parcellation of resting-state fMRI data (Yeo et al., 2011). These are shown in Figure 2. There are two important points to take away from Figure 2 that colour the results reported below. First, the functional networks frequently overlap one another. The social network overlaps considerably with the emotion network, particularly in the amygdala and ventromedial prefrontal cortex. It also shows convergence with situation-processing regions in the lateral anterior temporal lobes and inferior parietal cortices. Parts of left inferior prefrontal cortex are implicated in semantic control but also more generally in language comprehension, presumably as control is frequently needed to resolve verbal semantic ambiguities (Vitello & Rodd, 2015). Even the sensorimotor networks for vision and action overlap in the lateral occipitotemporal cortex, around motion-sensitive area V5/MT+ (Zeki, 2004). This overlap in functional networks reflects the fact that higher-order cognition involves the co-activation of a broad range of representations and processes. For example, understanding situations involving people may require us to simulate their mental states, leading to overlap between the situation and social networks. This cements the idea that different perspectives on concreteness effects are not mutually exclusive. It is possible, for example, that abstract concepts rely heavily on language representations and, as a consequence, recruit semantic control regions that resolve ambiguity in language. By understanding how different networks work in concert to support concrete and abstract comprehension, we gain insights into how meaning is constructed among multiple interacting systems.

**Figure 2:**
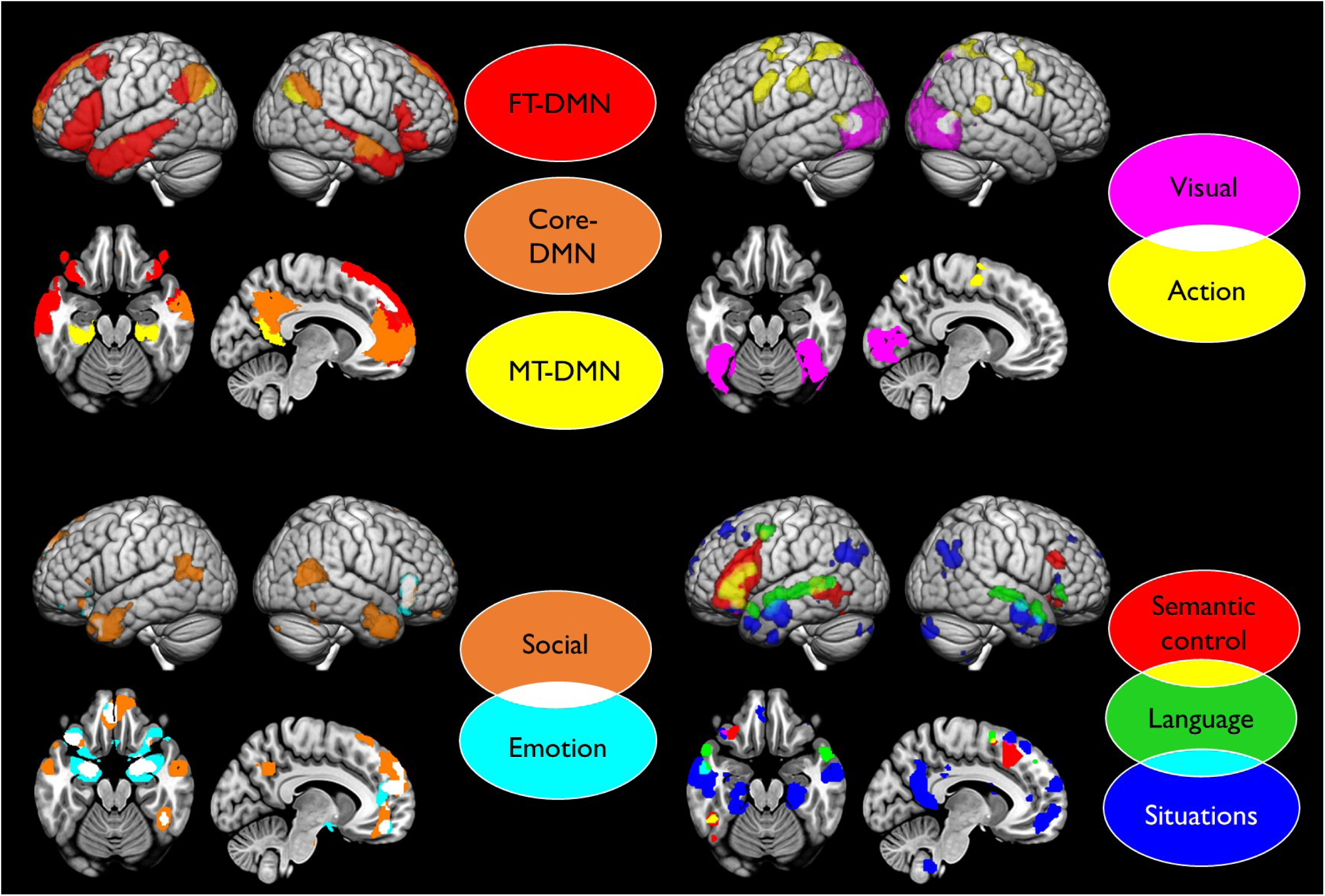
Functional networks used to interpret concreteness effects. *DMN = default mode network; FT = fronto-temporal; MT = medial temporal*.

The second critical point is that many of the functions under investigation align with sub-divisions of the DMN. As other researchers have noted, regions involved in social cognition, language processing and semantic control tend to fall within FT-DMN (particularly in the lateral temporal and lateral prefrontal cortices) but not MT-DMN (Andrews-Hanna et al., 2014; Smallwood et al., 2021). This could bias FT-DMN regions towards preferring abstract concepts. In contrast, the situation-processing network includes parts of all DMN sub-divisions, including lateral temporal regions of FT-DMN and medial temporal and retrospenial parts of MT-DMN. This may reflect the diverse types of processing that are relevant to different types of events and situations. For example, inter-personal interactions are important in many real-world situations, and these may engage social cognition regions in FT-DMN. In other situations, the physical environment may be more salient and these could engage scene processing regions in MT-DMN. The general alignment of the situation network with DMN fits with the view that a major function of the DMN is to construct situation models (Baldassano et al., 2018; Heidlmayr et al., 2020; Morales et al., 2022; Ranganath & Ritchey, 2012; Yeshurun et al., 2021). The association of the situation network with multiple DMN sub-divisions hints at the diverse types of processing called upon across situations and also raises the possibility that concrete and abstract specialisation might occur in different parts of this network. In contrast, there is no intersection between action and visual networks and the DMN. MT-DMN regions, particularly in the medial temporal lobe, are linked with the mental representation of visual scenes (Epstein & Baker, 2019). However, temporal lobe MT-DMN regions fall slightly anterior to those implicated in direct visual perception.

### Overall Effects of Concreteness

Activation Likelihood Estimation (ALE) analyses (Eickhoff et al., 2009, 2012) were used to identify regions that consistently showed positive (C>A) and negative (A>C) concreteness effects across the full set of 72 studies. Results are shown in Figure 3 (for peak co-ordinates, see Table 2). The most prominent A>C effects were observed in left inferior frontal gyrus (IFG) and anterior parts of the superior and middle temporal gyri (aSTG/MTG). C>A effects were most pronounced in the angular gyri (AG) and posterior parts of the left inferior temporal, fusiform and parahippocampal gyri (ITG/FG/PHG). Neighbouring regions of cortex sometimes showed opposing effects, particularly in posterior MTG/ITG and in the posterior cingulate and precuneus (PCC/PCu). In addition, while previous meta-analyses identified strongly left-lateralised effects, our more comprehensive analysis reveals that these effects are often mirrored in the right hemisphere, particularly in right aSTG/MTG (A>C), right PHG (C>A) and right AG (C>A). Thus, although language processing is traditionally thought of as a left-lateralised function, concrete and abstract words differentially engage bilateral brain networks, in line with the bilateral nature of many of the functional networks shown in Figure 2.

**Figure 3:**
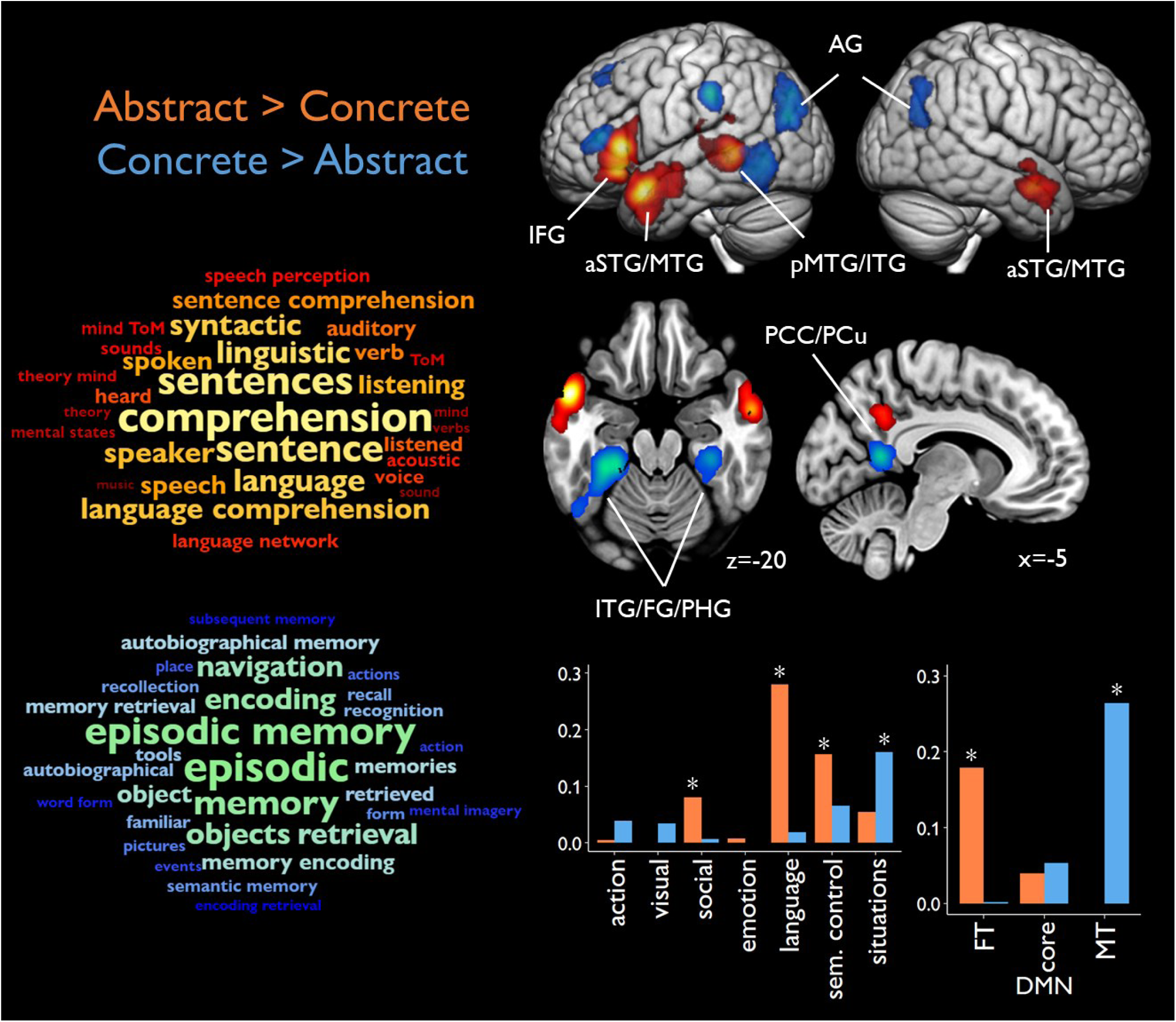
Meta-analysis of 72 studies of concreteness effects. Greater activation for abstract concepts (A>C) shown in hot colours and for concrete concepts (C>A) shown in cool colours. Results are shown at a voxel-level threshold of p<0.005, corrected for multiple comparisons at the cluster level. Wordclouds show Neurosynth terms most associated with A>C and C>A effects. Bar plots indicate Jaccard similarity between regions showing significant effects and the networks shown in Figure 2. * indicates above-chance similarity (FDR-corrected p < 0.05).

**Table 2:** Peak effect co-ordinates for the main analysis.

| Cluster | Regions | Volume<br>(mm<sup>3</sup>) | x | y | z | ALE value |
| --- | --- | --- | --- | --- | --- | --- |
| Abstract > Concrete |  |  |  |  |  |  |
| 1 | Left inferior frontal gyrus & lateral anterior temporal lobe | 18448 | -54 | 22 | 6 | 0.053 |
|  |  |  | -50 | 10 | -22 | 0.051 |
|  |  |  | -48 | 24 | -8 | 0.042 |
|  |  |  | -50 | 8 | -36 | 0.023 |
|  |  |  | -58 | -8 | -18 | 0.020 |
|  |  |  | -58 | -10 | -10 | 0.018 |
| 2 | Left posterior middle and superior temporal gyri | 7408 | -58 | -40 | 0 | 0.050 |
|  |  |  | -60 | -26 | 8 | 0.014 |
|  |  |  | -56 | -26 | 22 | 0.012 |
|  |  |  | -60 | -22 | 16 | 0.012 |
| 3 | Right lateral anterior temporal lobe | 5200 | 56 | 0 | -18 | 0.032 |
|  |  |  | 52 | 8 | -24 | 0.022 |
|  |  |  | 52 | -12 | -14 | 0.019 |
|  |  |  | 54 | -12 | -4 | 0.018 |
|  |  |  | 62 | -2 | -4 | 0.016 |
| 4 | Posterior cingulate & precuneus | 3328 | 2 | -54 | 30 | 0.025 |
|  |  |  | -10 | -52 | 36 | 0.022 |
|  |  |  | -4 | -40 | 32 | 0.014 |
| 5 | Left parietal operculum | 2216 | -48 | -38 | 22 | 0.022 |
| Concrete > Abstract |  |  |  |  |  |  |
| 1 | Left posterior middle and inferior temporal gyri, fusiform & parahippocampal gyrus | 14520 | -30 | -36 | -18 | 0.049 |
|  |  |  | -24 | -32 | -22 | 0.045 |
|  |  |  | -54 | -62 | -2 | 0.033 |
|  |  |  | -52 | -60 | -12 | 0.029 |
| 2 | Left angular gyrus & lateral occipital cortex | 9000 | -38 | -74 | 34 | 0.036 |
|  |  |  | -44 | -76 | 24 | 0.035 |
|  |  |  | -34 | -82 | 28 | 0.023 |
|  |  |  | -38 | -74 | 48 | 0.015 |
| 3 | Posterior cingulate & precuneus | 4800 | -8 | -54 | 10 | 0.040 |
|  |  |  | 8 | -54 | 10 | 0.021 |
|  |  |  | -14 | -50 | 26 | 0.014 |
| 4 | Right angular gyrus | 4424 | 46 | -66 | 24 | 0.030 |
|  |  |  | 44 | -60 | 40 | 0.021 |
|  |  |  | 48 | -68 | 46 | 0.017 |
| 5 | Left inferior frontal sulcus | 3368 | -42 | 32 | 12 | 0.034 |
| 6 | Left superior frontal gyrus | 3296 | -20 | 16 | 52 | 0.029 |
|  |  |  | -24 | 28 | 52 | 0.018 |
|  |  |  | -26 | 8 | 50 | 0.018 |
|  |  |  | -28 | 34 | 46 | 0.017 |
|  |  |  | -24 | 36 | 48 | 0.017 |
| 7 | Right parahippocampal gyrus | 2848 | 32 | -30 | -18 | 0.032 |
|  |  |  | 30 | -40 | -24 | 0.018 |
| 8 | Left supramarginal gyrus | 2416 | -60 | -30 | 38 | 0.044 |

The bar plots in Figure 3 show the Jaccard similarity between the regions showing concreteness effects and our functional networks of interest, with asterisks indicating where similarity is significantly greater than expected by chance (at FDR-corrected *p* < 0.05). These results support some of the predictions in Table 1. Abstract words preferentially activated the social network (particularly aSTG/MTG), language network (IFG and lateral temporal cortex) and semantic control network (left IFG and pMTG). Concrete word effects aligned with the situation network, primarily due to bilateral effects in PCC/PCu, PHG and AG. There was no significant relationship with the action network, perhaps because many studies did not include verbs in their stimuli (as we shall see, sentence stimuli are more likely to elicit C>A effects in this network). More surprisingly, C>A effects were not significantly aligned with the visual network. Concrete word effects in the left pITG partially overlapped with lateral occipitotemporal regions involved in visual object recognition, though there was no C>A effect in the homologous right-hemisphere region. There were also areas of C>A effects in ventral occipitotemporal cortex bilaterally; however, the strongest effects were anterior to the regions implicated in visual processing. Overall, these limited overlaps did not amount to a statistically significant association. As this result was unexpected, we repeated the analysis using alternative definitions of visual cortex based on object recognition functional localisers and retinotopic mapping (Rosenke et al., 2021; L. Wang et al., 2015). These alternative visual networks also showed no significant association with C>A effects (see Supplementary Information for details).

When we considered DMN sub-networks, we saw a striking dissociation between concrete and abstract concepts. Greater activation for abstract words was strongly associated with FT-DMN regions while greater activation for concrete words overlapped significantly with MT-DMN regions.

This dissociation is clearest in the anterior temporal cortices, where lateral FT-DMN regions were more active for abstract language while ventromedial MT-DMN regions preferred concrete language (for a similar result, see (Hoffman et al., 2015)). It is also evident in left IFG (A>C and FT-DMN), and in the retrospenial cortex and posterior portions of the AG (C>A and MT-DMN). These results have two important implications. First, although DMN is most associated with tracking semantic content in complex narratives (Baldassano et al., 2018; Heidlmayr et al., 2020; Morales et al., 2022), our results show that differential DMN engagement is widely elicited by more simple language stimuli (single words and sentences). Second, they indicate that the type of content that engages this system varies systematically along a social-spatial axis.

Finally, the wordclouds in Figure 3 visualise the terms in the neuroimaging literature that are most strongly associated with regions showing C>A and A>C effects. These were generated by submitting C>A and A>C ALE maps to the decoding function in the Neurosynth meta-analytic database (Rubin et al., 2017; Yarkoni et al., 2011) (see Method for details). This automated decoding approach supports the picture already seen. It confirms that regions preferentially activated by abstract language are associated in the literature with theory of mind, mental states and language processing, particularly at the sentence level. Conversely, the terms associated with concrete regions include aspects of visual and action processing (e.g., tools, pictures, mental imagery) but are dominated by episodic and autobiographical memory, which are core MT-DMN functions. To ensure that these effects did not stem from task instructions, we identified 7 studies in the meta-analysis where participants were asked to either encode or retrieve items from episodic memory and re-ran the analysis excluding them. The exclusion of these studies did not change the results (see Supplementary Figure 1), indicating that these effects do not stem from explicit demands in episodic memory. One possibility is that concrete words automatically trigger retrieval of autobiographical or episodic memories, because they bring to mind specific situations (Binder, 2016). Alternatively, it could be that tasks typically used to probe episodic/autobiographical memory rely on similar processes to those involved in comprehending concrete language (e.g., the mental representation of visuospatial contexts).

### Influence of study characteristics

We next tested how concreteness effects varied as a function of stimulus and task. Figure 4 shows contrasts of Word vs. Sentence studies, conducted separately for A>C and C>A effects. Red regions in the left-hand panel are those where A>C effects were more likely to be observed for studies that presented sentences (or multi-word phrases). These regions primarily overlapped with the social and situation networks and included the lateral anterior temporal lobes bilaterally and left temporoparietal junction. These findings support the idea that abstract sentences evoke richer representations of situations than individual abstract words, and that these situations frequently include social elements. Yellow regions were more likely to show A>C effects in studies that presented single words. These effects were concentrated in left IFG and pMTG, which are major constituents of the semantic control network (Jackson, 2021). These results are consistent with the view that the cognitive control demands of processing abstract words are maximal when no context is available to constrain their interpretation.

**Figure 4:**
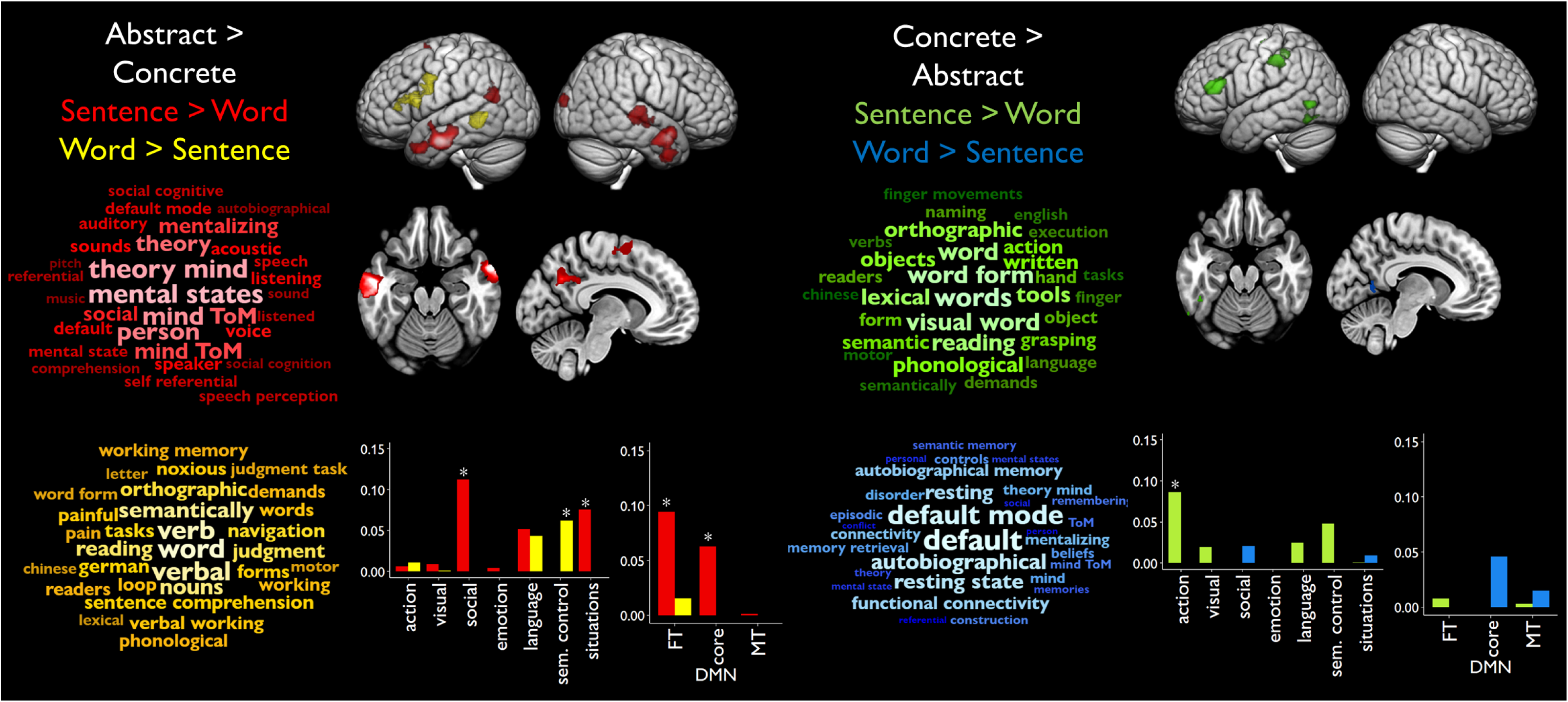
Effects of stimulus type. Figure shows regions where A>C (left side) and C>A (right side) effects were more likely for Sentence vs. Word studies. Wordclouds show Neurosynth terms most associated with regions showing effects. Bar plots indicate Jaccard similarity between regions showing significant effects and the networks shown in Figure 2. * indicates above-chance similarity (FDR-corrected p < 0.05).

The right-hand panel of Figure 4 shows stimulus influences on concrete word activation. Green regions show where C>A effects were more likely for Sentence studies. These effects were observed mainly in parts of the action network, specifically left lateral occipitotemporal cortex and supramarginal gyrus. This engagement suggests simulation of sentence content, since concrete sentences invariably describe physical actions. In contrast, Word studies typically present concrete nouns rather than verbs, which might not be expected to engage simulations of actions. A region of the left inferior frontal sulcus was also more likely to show C>A activation in Sentence studies. Greater likelihood of C>A activation in Word studies was limited to a small number of voxels in ventral precuneus.

Figure 5 presents analysis of studies split by task type. For A>C effects, a number of regions were more likely to show abstract word preferences when participants made explicit semantic judgements about the stimuli (or opposed to non-semantic tasks or passive reading/listening). These regions overlapped primarily with the language network, suggesting that deeper semantic processing for abstract concepts relies on the language system. No regions were more likely to show A>C effects in Non-semantic studies. Semantic studies were more likely to find C>A activation in a distributed set of regions that primarily coincided with the situation network. These included parts of AG, PCu and PHG. Greater activation in Non-semantic studies was confined to a single cluster in the left pre-central gyrus, a region primarily associated with action and motor processing. These effects might reflect a tendency to focus on motoric aspects of concrete concepts during shallower processing (though these effects did not show significant correspondence with the Action network as a whole). Finally, when comparing across panels in Figure 5, it is clear that the dissociation in concreteness effects across the DMN was more likely to occur when participants made explicit semantic judgements. This suggests the association of the two classes with different DMN sub-divisions is related to deep processing of the conceptual significance of these concepts.

**Figure 5:**
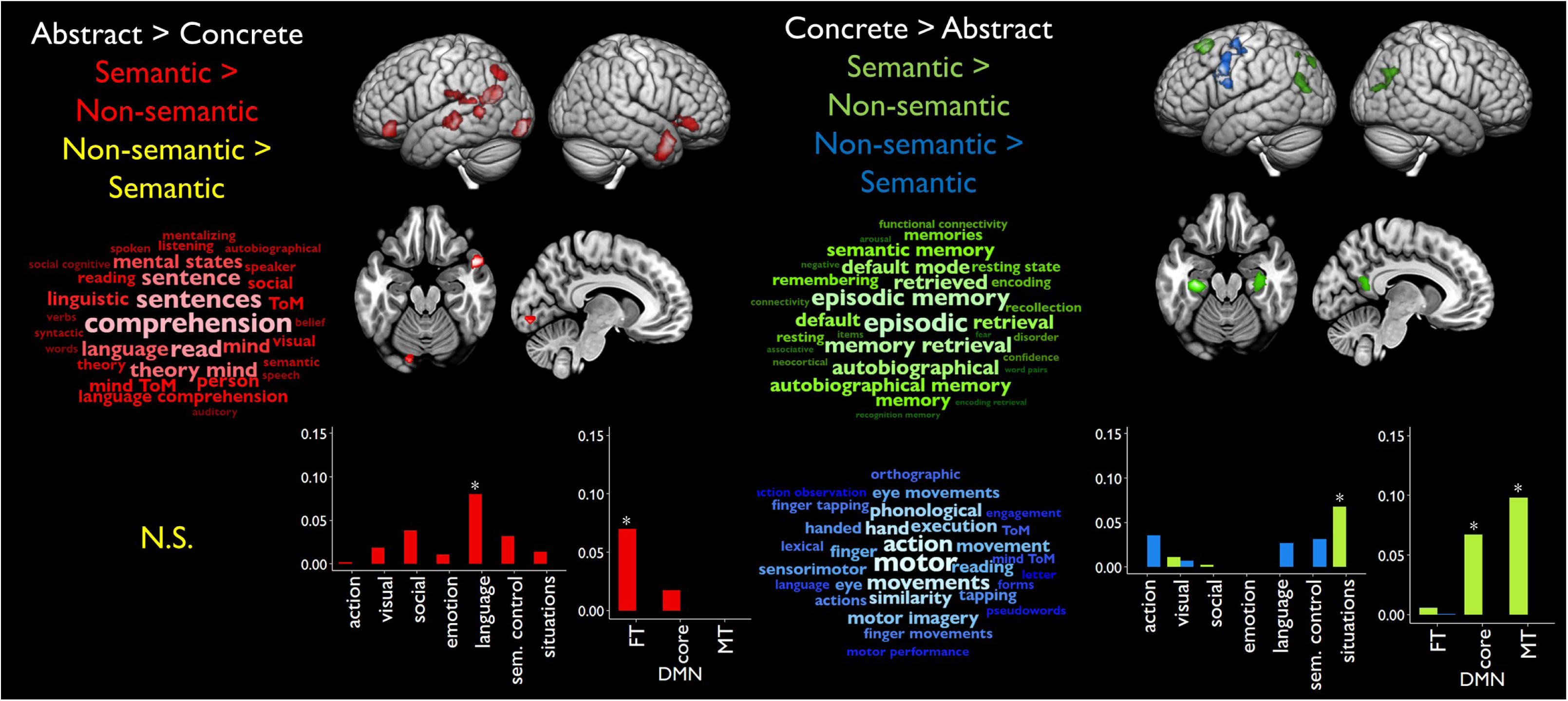
Effects of task type. Figure shows regions where A>C (left side) and C>A (right side) effects were more likely for Semantics vs. Non-semantic studies. Wordclouds show Neurosynth terms most associated with regions showing effects. Bar plots indicate Jaccard similarity between regions showing significant effects and the networks shown in Figure 2. * indicates above-chance similarity (FDR-corrected p < 0.05). N.S. = nothing significant.

## Discussion

The distinction between concrete and abstract concepts is fundamental to language and semantic cognition, with a range of theories predicting specific differences in how these classes of concept are processed in the brain. We tested these predictions in the largest meta-analysis of the literature to date. When aggregating across 72 neuroimaging studies, we found widespread activation differences between concrete and abstract language, falling within a range of functional networks that support different aspects of conceptual processing. In particular, there was a systematic association of concrete and abstract activation with different subdivisions of the DMN. Effects were most widespread in the left hemisphere though in some cases these were mirrored in the right, in line with other findings that conceptual processing in language tasks engages bilateral neural networks (Patel et al., 2023; Yu et al., 2024). We will now consider the implications of our results for the concreteness theories outlined earlier and for the role of DMN regions in supporting mental representation, before considering some limitations of our approach.

Many of the predictions relating to abstract language activation were supported by our analyses (see Table 1). Abstract language preferentially activated the social network and this effect was more likely when participants processed sentences rather than single words. Sentences are presumably able to elicit richer representations of social interactions whereas without this context, the social significance of a single abstract word may be fairly impoverished (Barsalou & Wiemer-Hastings, 2005; Davis et al., 2020). We found no evidence, however, that abstract concepts preferentially activate emotion-processing regions, contrary to the Affective Embodiment hypothesis (Kousta et al., 2011; Lenci et al., 2018; Newcombe et al., 2012; Vigliocco et al., 2014). Cross-linguistic analyses have found that while abstract words are, on average, more strongly emotionally valenced than concrete words, this effect is driven by a small subset of abstract words that have strong emotional connotations (Winter, 2023). If these highly emotional words form only a small proportion of the abstract words used in most studies, then reliable effects in emotion systems would not be expected. Thus, our results caution against considering affective embodiment as a general feature of abstract language comprehension (unlike social processing), though it may be important for particular subsets of words.

Abstract concepts engaged the language comprehension network to a greater degree than concrete, as would be expected if language is a more central modality for the acquisition and use of these concepts (Borghi et al., 2019; Vigliocco et al., 2009; Villani et al., 2019). This effect was more likely to occur when people made semantic judgements about stimuli (cf. non-semantic or passive tasks). This result could call into question the claim that linguistic representations are most useful for superficial processing (Barsalou et al., 2008; Connell, 2019), though such an argument depends on the difficulty and depth of the processing required by the semantic studies. Many of the semantic tasks involved straightforward processing like synonym matching or detection of semantic violations, which could potentially be supported by the activation of linguistic associations. Abstract concepts also relied more heavily on semantic control regions, supporting the claim that the complex, variable meanings of these words require more executive regulation (Bechtold et al., 2021; Cousins et al., 2016; Davis et al., 2020; Hoffman, 2016; Hoffman et al., 2010, 2015). This effect was particularly prominent for single-word processing, in line with the claim that when abstract words are placed in context, the ambiguity surrounding their meaning is reduced (Davis et al., 2020; Hoffman et al., 2010, 2015).

Embodied semantics theories (Barsalou, 1999, 2008; Binder, 2016; Binder & Desai, 2011; Connell et al., 2018; Gallese & Lakoff, 2005; Glenberg & Robertson, 2000; Lambon Ralph et al., 2017; Martin, 2016; Pulvermüller, 2013) predict that concrete language should produce greater activation in the brain’s sensorimotor systems, as the process of comprehending them involves simulation of relevant perceptual and motor experiences. We found mixed evidence for this prediction. Preferential activation for concrete stimuli did not overlap with the action network in the main meta-analysis, but concrete effects in this network were more likely for sentence stimuli. Thus, action-processing regions appear to be preferentially activated when people process concrete sentences (which almost invariably describe physical movements or actions) compared with abstract sentences. The lack of an effect in the main analysis is likely due to the preponderance of single-word studies, which present participants with nouns more frequently than verbs.

We found no significant overlap between concrete-preferring regions and the visual system, neither in the main analysis or when using various alternative ways of defining visual regions. This is surprising because concreteness is defined as the degree to which concepts are experienced through the senses and visual experience seems to be the dominant sensory modality for most concrete words (Lynott et al., 2020). The lack of an effect in visual regions is hard to reconcile with strong embodiment theories which hold that activation of sensorimotor cortex is an automatic part of all language comprehension (Gallese & Lakoff, 2005; Glenberg & Robertson, 2000). Our results are more compatible with weaker embodiment positions which hold that sensorimotor simulations are engaged to varying degrees depending on context and task (Barsalou et al., 2008; Binder, 2016; Binder & Desai, 2011; Lambon Ralph et al., 2017; Zwaan, 2014). This class of theories sees simulation as one aspect of conceptual representation that can be drawn upon where appropriate, while in other situations people rely more on linguistic or modality-independent semantic representations. Studies of concreteness tend not to ask participants to explicitly engage in mental imagery or bring to mind specific perceptual properties, because of the difficulty of applying the same instructions to abstract concepts. Thus, in the studies we have analysed, people are rarely required to generate strong visual simulations and this may explain the lack of effects in visual cortex. In contrast, visual and action regions are regularly recruited when people are specifically instructed to focus on the perceptual qualities of concrete words (Fernandino et al., 2016; Martin et al., 1995; Thompson-Schill et al., 1999; van Dam et al., 2012).

In addition to providing robust empirical assessment of long-standing claims about concreteness effects, our study provides a rich source of data to assess recent theories about the role of situation models in processing these concept classes (Barsalou et al., 2018; Barsalou & Wiemer-Hastings, 2005; Binder, 2016; Davis et al., 2020; McRae et al., 2018). Overall, concrete-preferring regions overlapped significantly with regions implicated in the processing of events and situations and this effect was strongest when people made semantic decisions. These results support claim that concrete words more readily evoke rich representations of specific situations (Binder, 2016; Schwanenflugel et al., 1988). Preferential activation for abstract words in the situation network was more likely when people processed sentences, suggesting that more abstract language can also elicit situation models when sufficient context is available. Importantly, concrete and abstract words engaged different parts of the situation processing network. Overlap for concrete words occurred primarily in retrospenial and medial temporal cortex while abstract effects clustered in the lateral anterior temporal lobes. This apparent divergence in situational processing is reflected in the clear dissociations we found within DMN sub-divisions, along a social-spatial axis. Abstract stimuli preferentially activated FT-DMN regions involved in social, language and semantic cognition while concrete stimuli engaged MT-DMN regions attuned to processing scenes and spatial contexts.

There is an emerging consensus that a general function of the DMN is to generate simulations of real or imagined events and contexts, variously termed situation models (Fernandino & Binder, 2024; Ranganath & Ritchey, 2012; Yeshurun et al., 2021), schemas (Gilboa & Marlatte, 2017), perceptually-decoupled cognition (Smallwood et al., 2013), mental time travel (Addis, 2020) or simply imagination (Andrews-Hanna & Grilli, 2021). In particular, Andrews-Hanna and Grilli (2021) propose that this general DMN function can be decomposed into two sub-systems they term the “mind’s eye” (MT-DMN), which represents visuospatial and external aspects of situations, and the “mind’s mind” (FT-DMN), which represents conceptual and internally-focused aspects. Previous studies have found that these systems are preferentially attuned to perceptual/spatial vs. thematic/conceptual aspects of autobiographical memories (D’Argembeau et al., 2014; Gurguryan & Sheldon, 2019), to memories of places vs. people (Silson et al., 2019) and to the vividness vs. emotional salience of imagined events (Lee et al., 2021).

Our data provide a powerful new source of evidence for this hypothesis by demonstrating that concrete and abstract language differentially engages the mind’s eye and mind’s mind. While a previous study has shown a similar pattern during story reading (Tamir et al., 2016), our meta-analysis shows that single sentences and even individual words are sufficient to drive selective responses in these DMN sub-systems, presumably because they spontaneously bring to mind models of relevant situations. Real-life situations involve a complex interplay of external and internally-focused elements, just as natural language uses a mixture of concrete and abstract words. Thus, natural situation models frequently need to integrate internal/social and external/spatial elements, explaining why the DMN frequently emerges as a single coherent network. In contrast, the studies in our meta-analysis have experimentally isolated activity relating to concrete and abstract aspects of thought. In so doing, we can begin to identify the underlying composition of the DMN simulation engine.

It is worth considering some alternative explanations for concreteness effects in the DMN. First, in line with its traditional characterisation as a task-negative network, DMN activation is known to negatively correlate with task difficulty (Mckiernan et al., 2003). Abstract language is usually more difficult to process than concrete (Schwanenflugel et al., 1988), which might lead to generic difficulty-related C>A effects in these regions. However, as we observed both C>A *and* A>C effects in parts of the DMN that show difficulty-related effects (e.g., PCC/PCu), this does not provide a straightforward explanation of our results. Second, MT-DMN regions are frequently implicated in episodic memory recall, raising the possibility that concrete concepts trigger associations with specific past experiences in a way that abstract concepts do not. While this is possible, it is more likely that similarities in activation arise from episodic memory tasks drawing on the mind’s eye simulation system. MT-DMN is activated by imagining future events as well as remembering past ones (Addis, 2020; Mullally & Maguire, 2014), suggesting a more general role in simulating both experienced and imagined events.

In this meta-analysis, we aimed to test the major established neurocognitive theories of concreteness. There are, however, some recent perspectives which we did not test explicitly. For example, we did not test recent proposals that comprehending abstract words entails greater activation of mouth motor cortex (Borghi et al., 2019) or that abstract concepts are particularly associated with internal bodily sensations (Connell et al., 2018). In addition, the meta-analytic approach identifies broad trends in activation that are consistent across a range of studies, tasks and stimuli. It is less sensitive to effects that may occur for specific subsets of concept within the broader concrete and abstract domains. For example, we did not find abstract word activation in emotion processing regions as has been proposed by others (Vigliocco et al., 2014) and this may be because emotions are only salient for a subset of abstract concepts that are not strongly sampled in most studies. There is a long tradition of investigating neural specialisation for categories of concrete concept, such as animals, plants and human-made artefacts (Chouinard & Goodale, 2010; J. T. Devlin et al., 2002). More recently, researchers have attempted to establish a similar taxonomy for abstract concepts, identifying sets of words that refer to social concepts, emotions, mental states and magnitudes amongst others (Conca et al., 2021; Troche et al., 2014).

This new focus on distinct categories of concept is a reminder that concreteness itself is a broad and multi-faceted construct. Unpicking the contributions of neural systems to specific concepts will require more nuanced methods than the uni-dimensional approach to concreteness that has dominated the field thus far. Some studies have moved in this direction, quantifying the contributions of specific sensorimotor domains (e.g., shape, colour, sound) as well as more intangible social, emotional, and cognitive aspects of experience to specific concepts (Binder et al., 2016; Connell et al., 2018; Hoffman & Lambon Ralph, 2013; Lynott et al., 2020; Troche et al., 2014; Villani et al., 2019). These studies have confirmed that concreteness is a fundamental organising principle, with concrete and abstract concepts strongly differentiated across many domains of experience. However, characterizing concepts at this finer-grained level allows for a more precise dissection of the neural systems underlying semantic representation. This can include identifying activation that correlates with specific domains of experience (Fernandino et al., 2016) and using multivariate approaches to identify regions whose representational structure is determined by experiential attributes (Tong et al., 2022; X. Wang et al., 2018).

These developments notwithstanding, the present study indicates that the broad classes of concrete and abstract concept vary reliably and systematically in their engagement of different neural networks. Our findings are well accounted for by two tenets of semantic cognition: 1. that semantic representations are partly embodied in modality-specific neural systems that process different aspects of our experiences (Barsalou, 1999, 2008; Binder, 2016; Binder & Desai, 2011; Connell et al., 2018; Gallese & Lakoff, 2005; Glenberg & Robertson, 2000; Lambon Ralph et al., 2017; Martin, 2016; Pulvermüller, 2013) and 2. that semantic activation is shaped by control processes, with the need for these varying with concept type and level of contextual support (Giallanza et al., 2023; Hoffman et al., 2010, 2015; Jackson, 2021; Lambon Ralph et al., 2017). Moreover, our analysis reveals that different subnetworks within the DMN are engaged by concrete vs. abstract language. We propose that this variation reflects differences in the types of situation models they elicit. Concrete language evoke situations in which representations of external space are dominant, while abstract language evokes situations in which social and linguistic features come to the fore.

## Acknowledgements

The research was supported by a BBSRC grant (BB/T004444/1). For the purpose of open access, the author has applied a Creative Commons Attribution (CC BY) licence to any Author Accepted Manuscript version arising from this submission.

## Supplementary Materials

**Supplementary Table 1:**
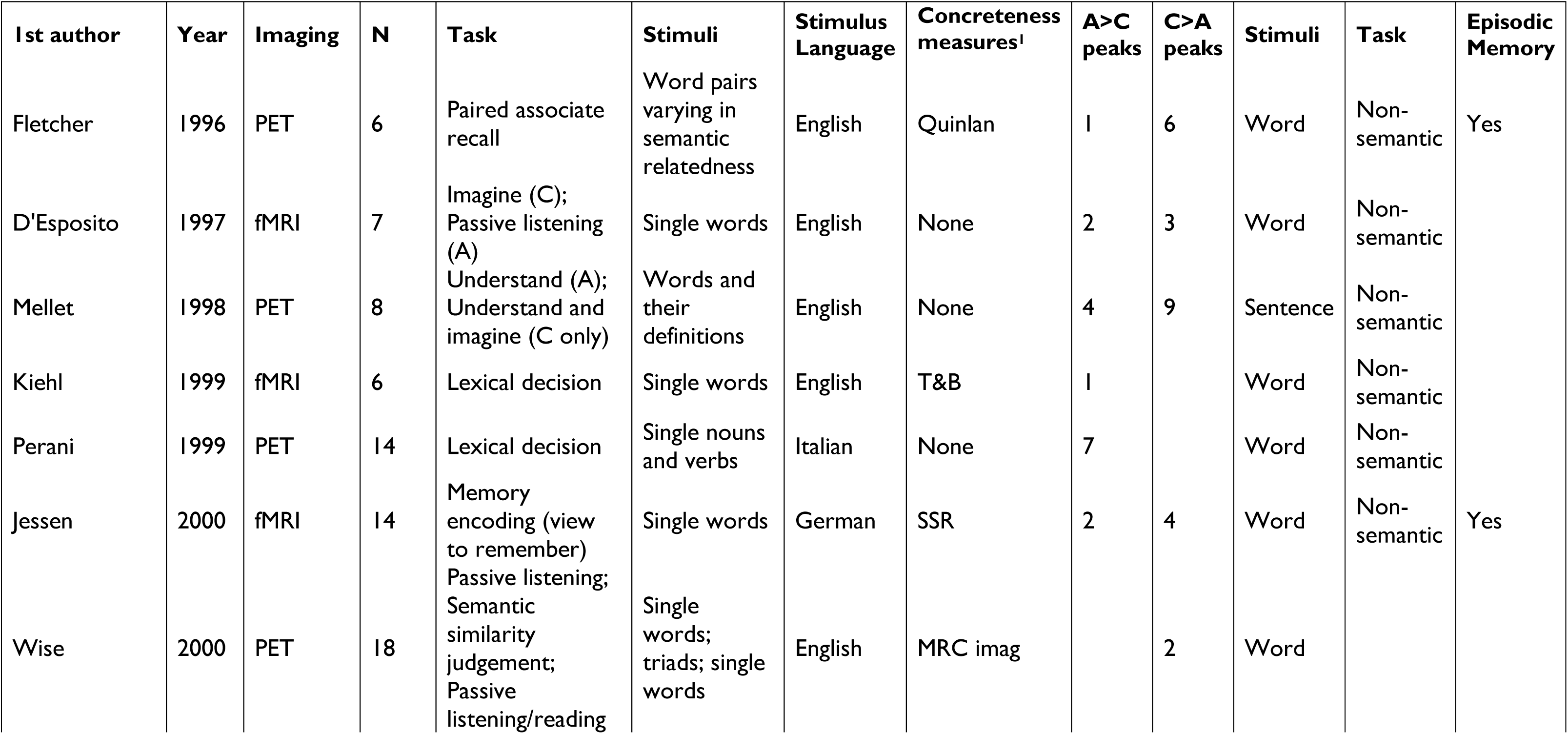
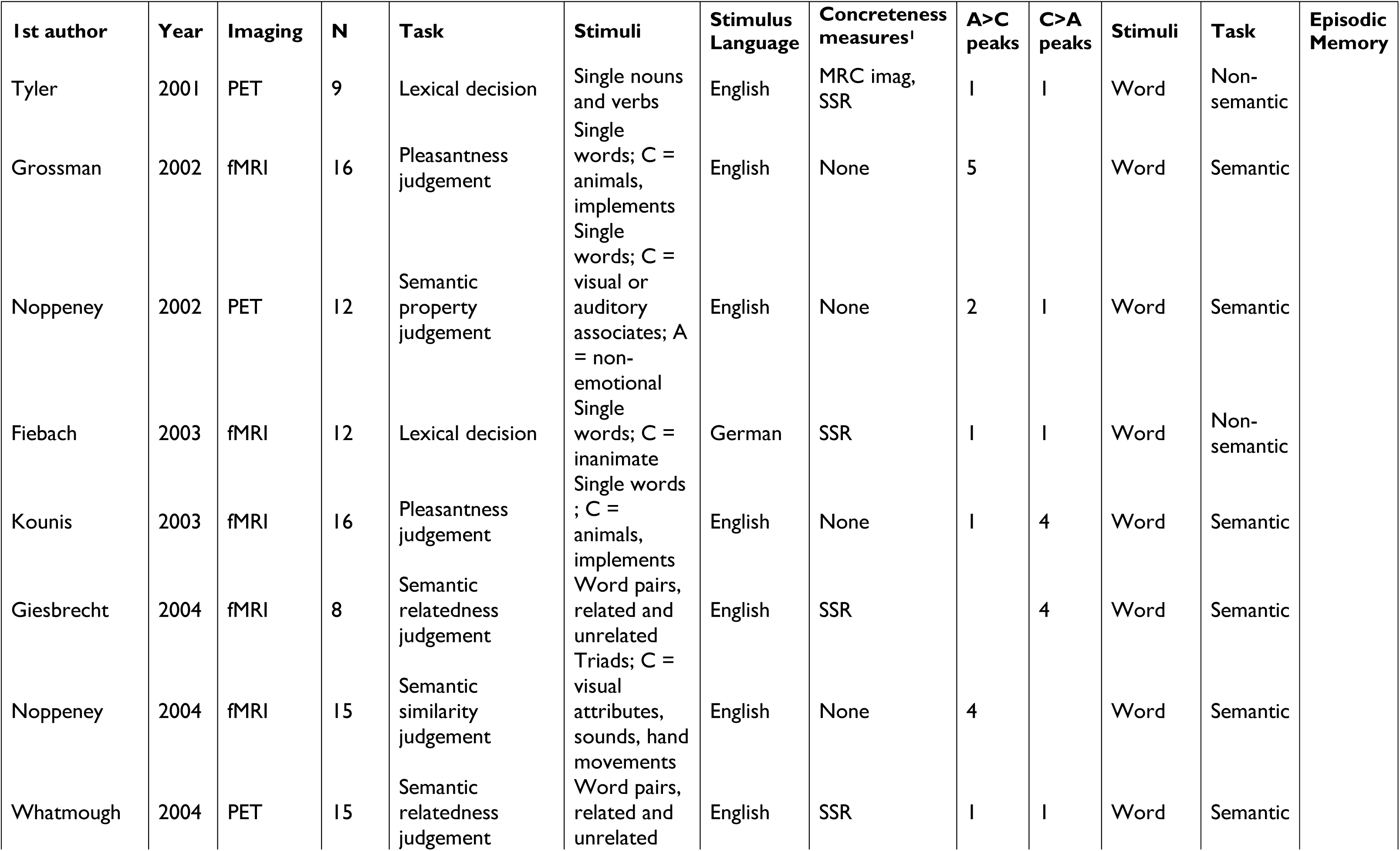
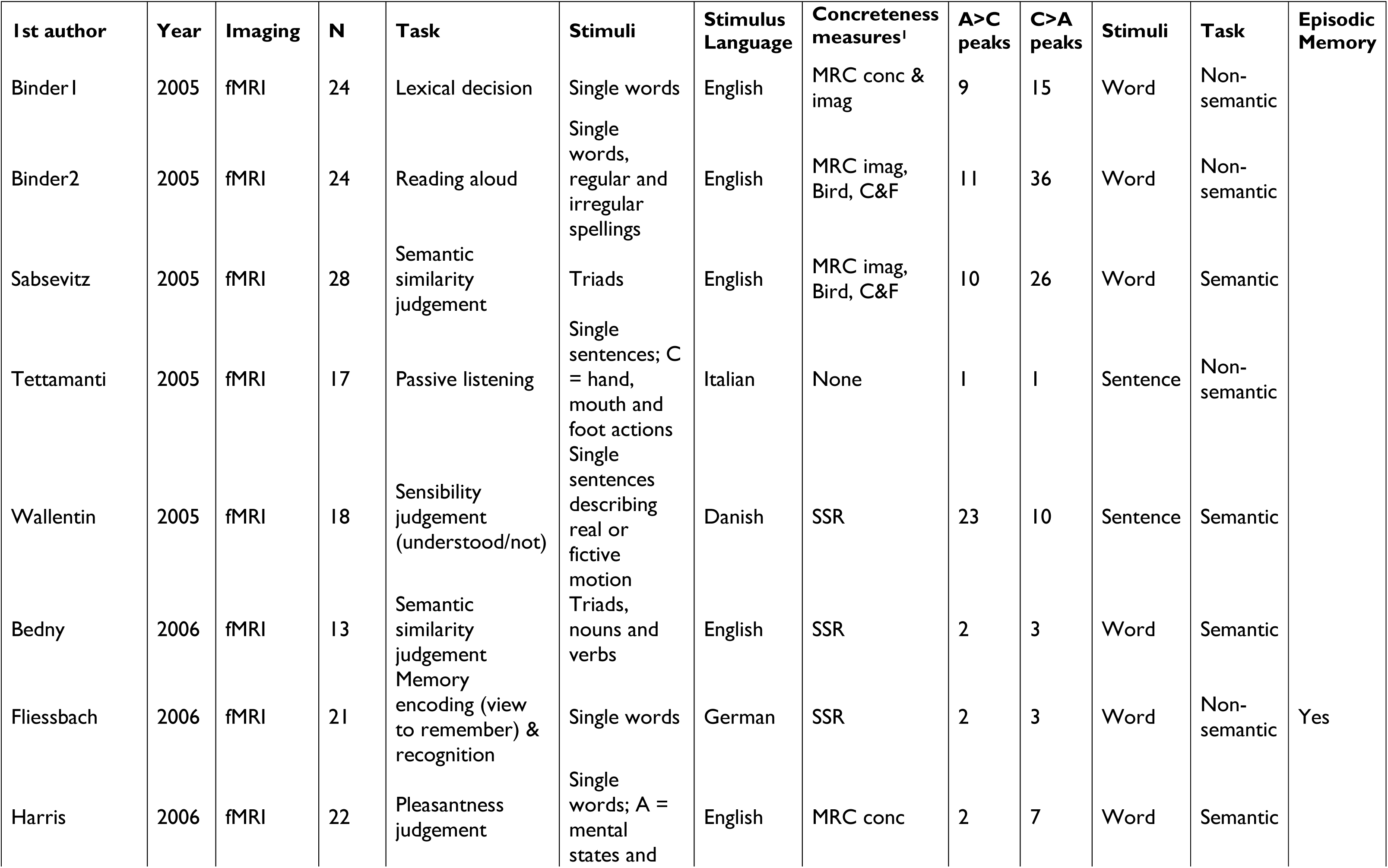
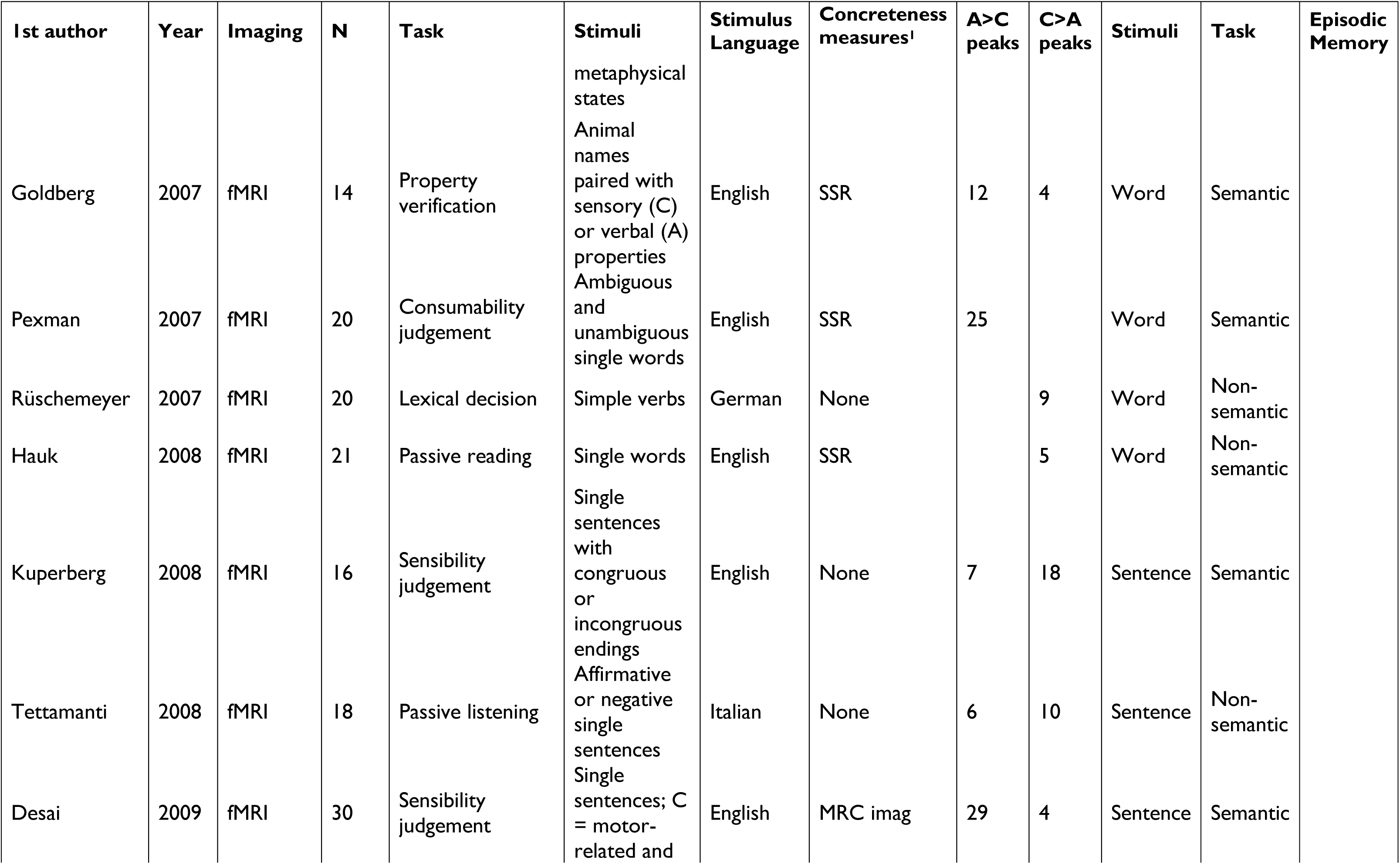
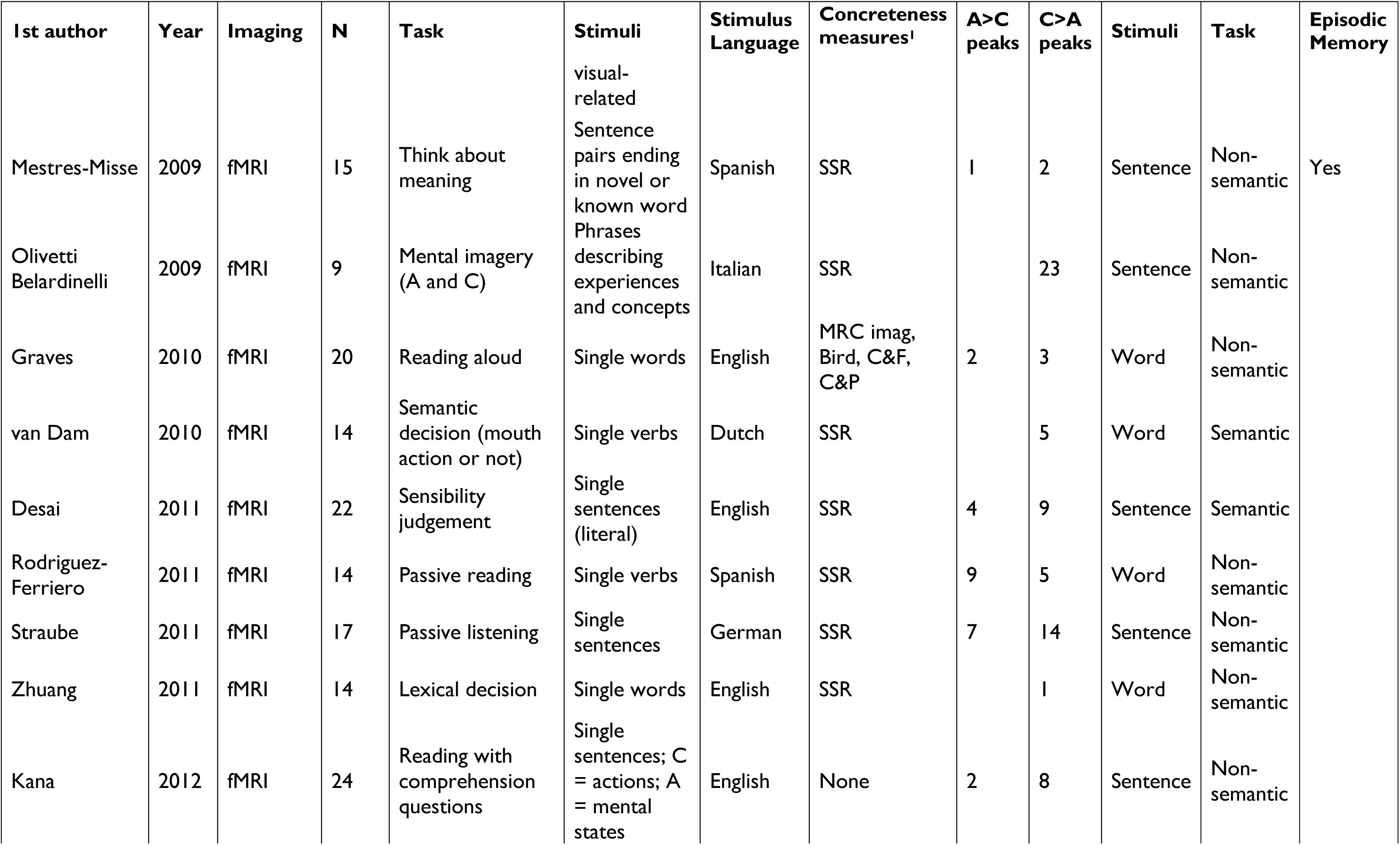
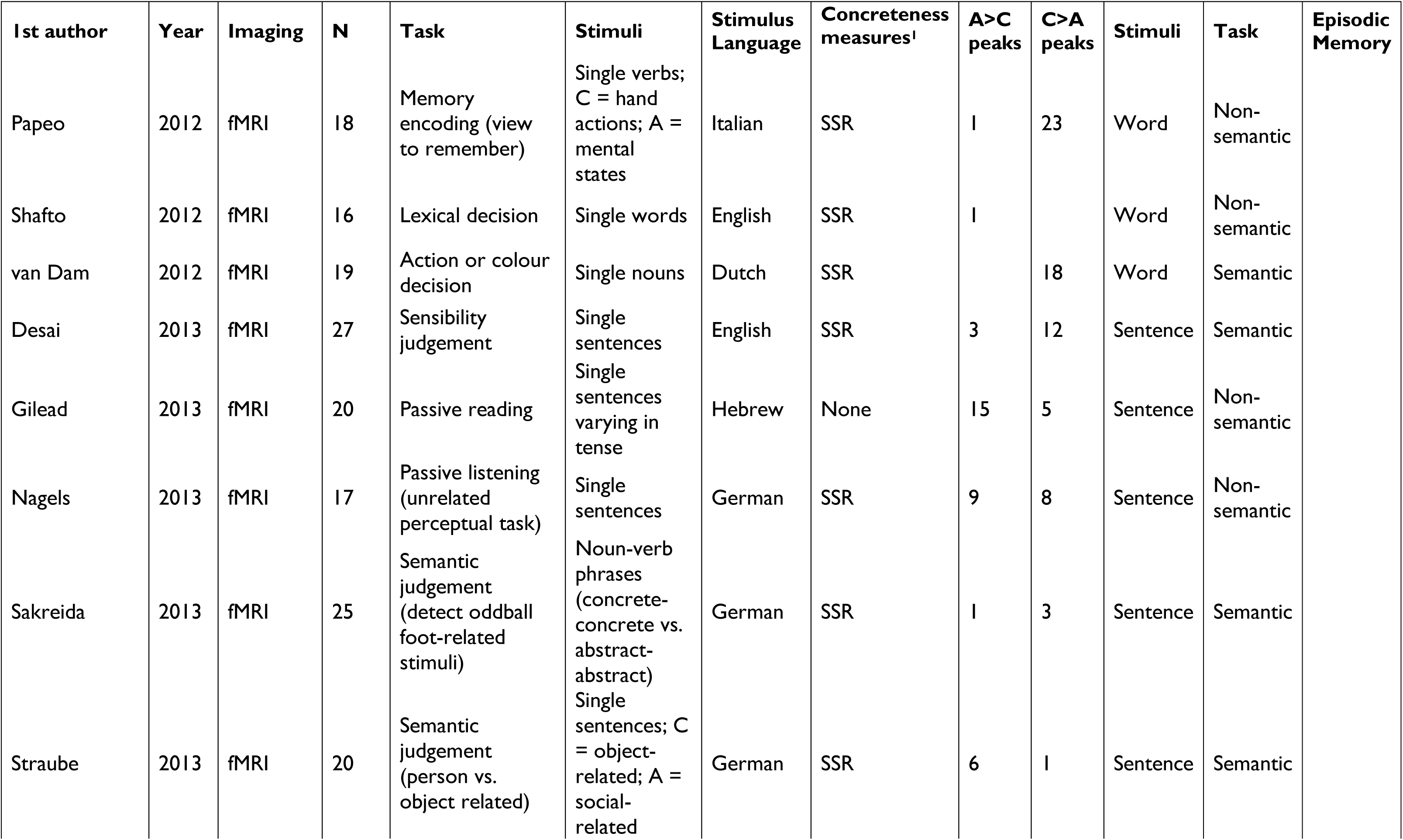
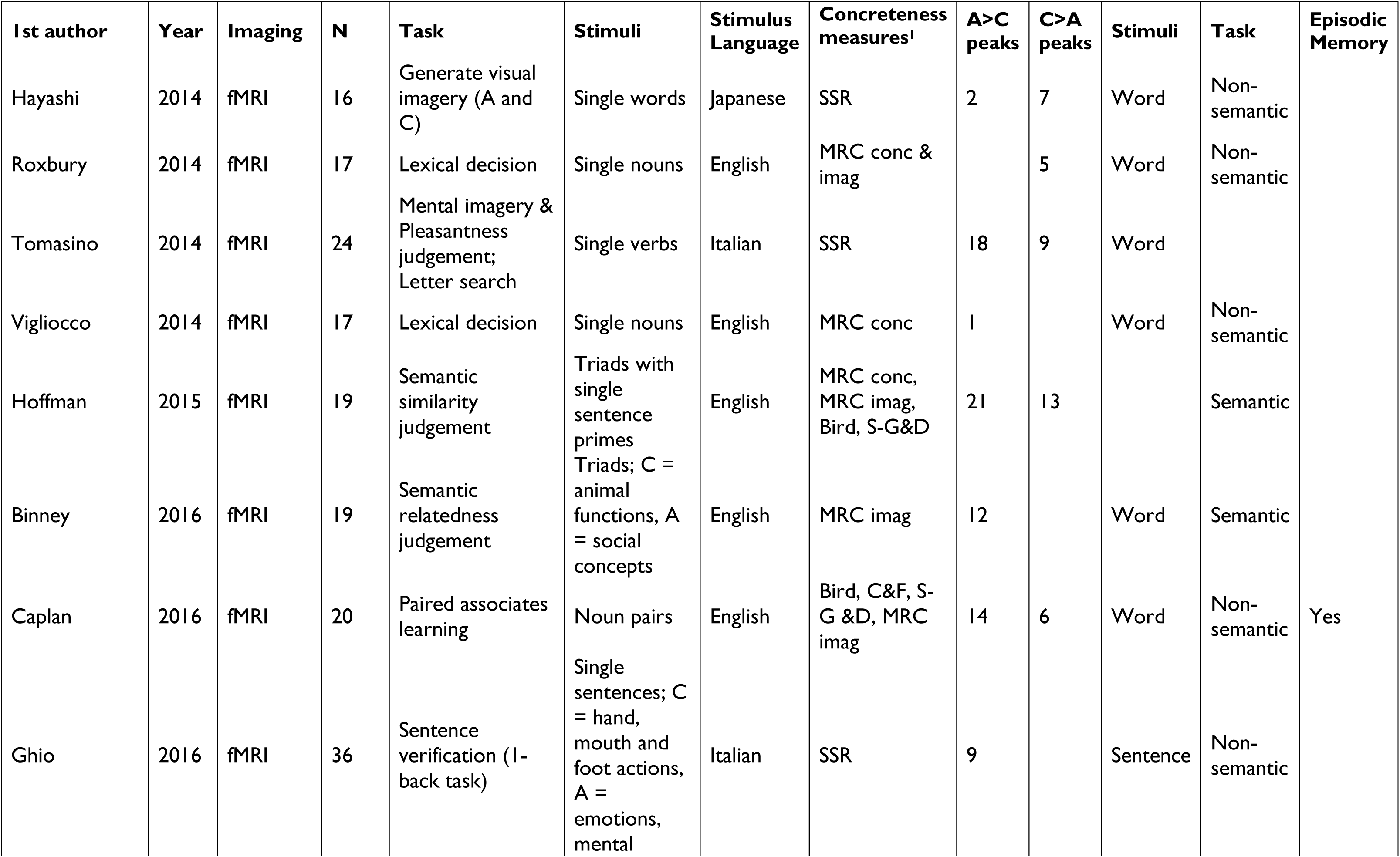
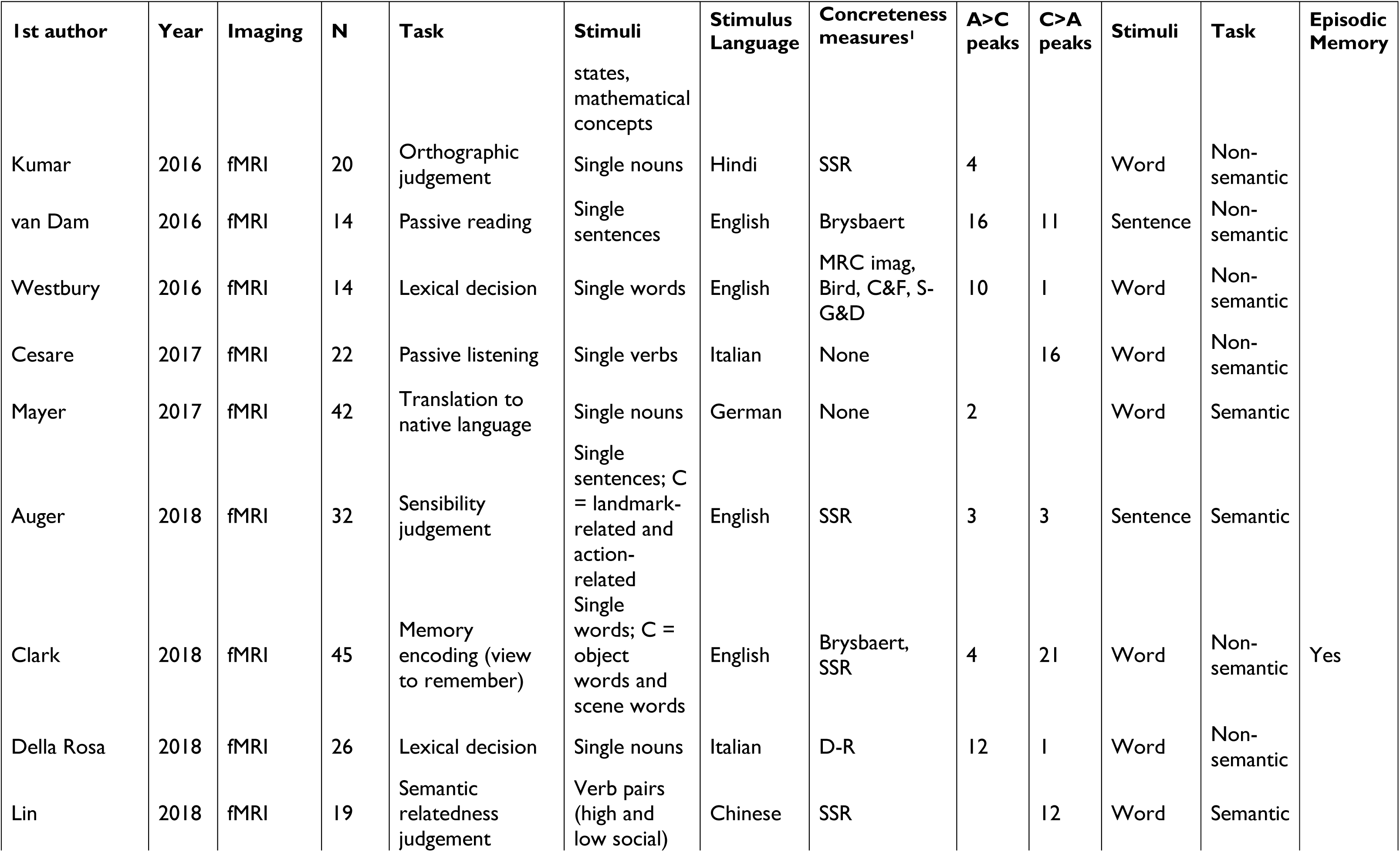
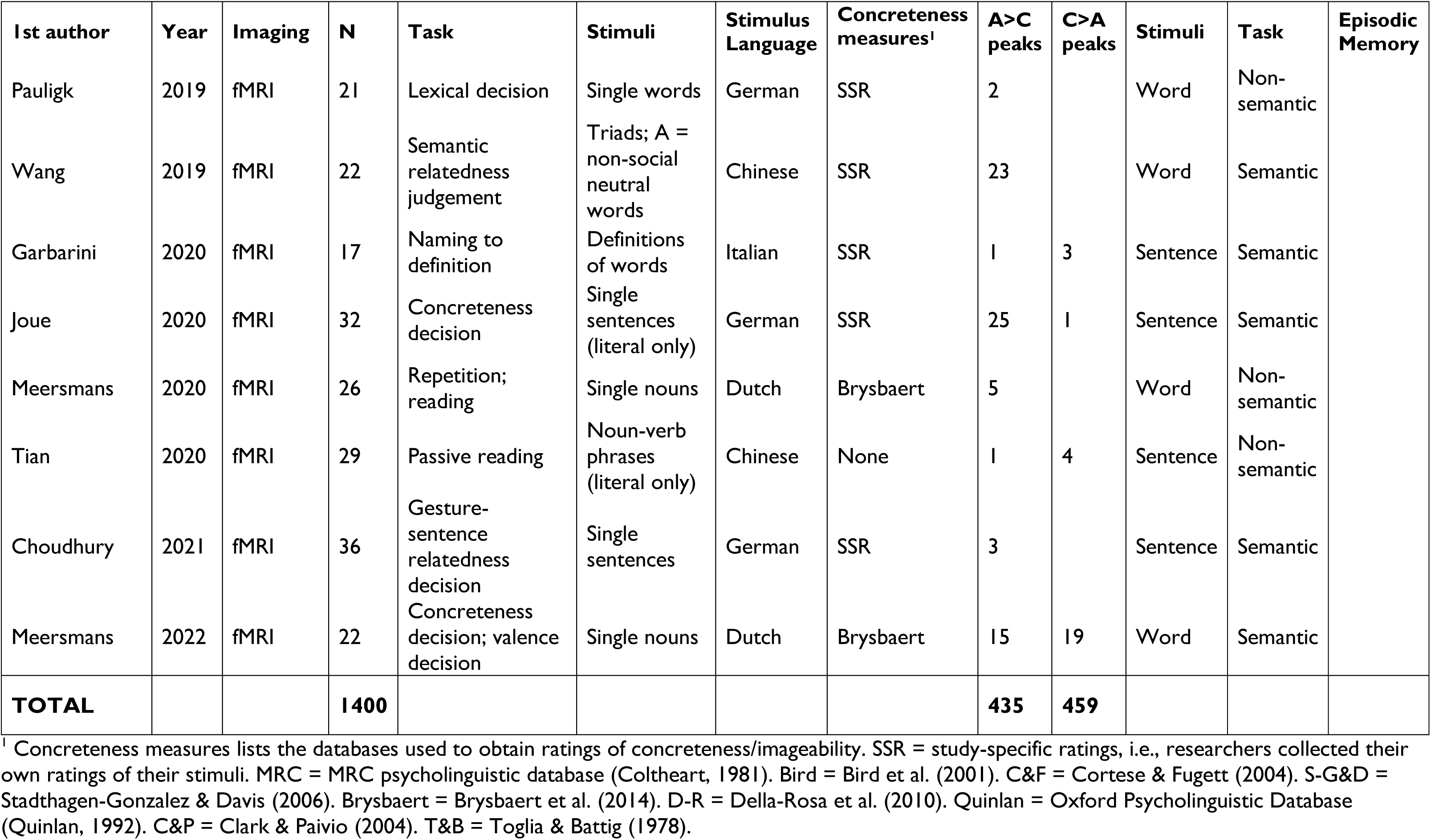
Details of all studies included in the meta-analysis.

### Supplementary Analysis: Alternative definitions of the visual system

The main analysis did not identify significant overlap between the visual system and regions showing preferential activation for concrete concepts. As this result was unexpected, we tested for overlap with three alternative visual networks defined in different ways.

1. *Neurosynth “ventral visual” map*. The overall “visual” network taken from the Neurosynth database includes large areas of parietal cortex that form the dorsal visual stream or “where/how” pathway. These regions might be less important for mental imagery of concepts than the ventral stream or “what” pathway, which is specifically implicated in object recognition.
2. *Rosenke et al. functional atlas*. This is a functional atlas of visual regions in the occipitotemporal cortex. It was generated by presenting a range of functional localiser tasks to 20 participants, including retinotopic mapping, motion vs. static stimuli and image presentation to identify category-selective regions (faces, bodies, places and characters).
3. *Wang et al. functional atlas*. This functional atlas provides visual topographic regions based on functional tasks performed in 53 participants. The protocol included retinotopic mapping and memory-guided saccade mapping but, unlike Rosenke et al., did not feature object recognition.

The three alternative visual networks are plotted alongside the C>A effects in the figure below. The first two networks show a small amount of overlap with concreteness effects in the left fusiform/inferior temporal gyrus, but they largely identify distinct areas. In no case did the Jaccard similarity exceed chance level (all uncorrected *p* > 0.05).

**Supplementary Figure 1:**
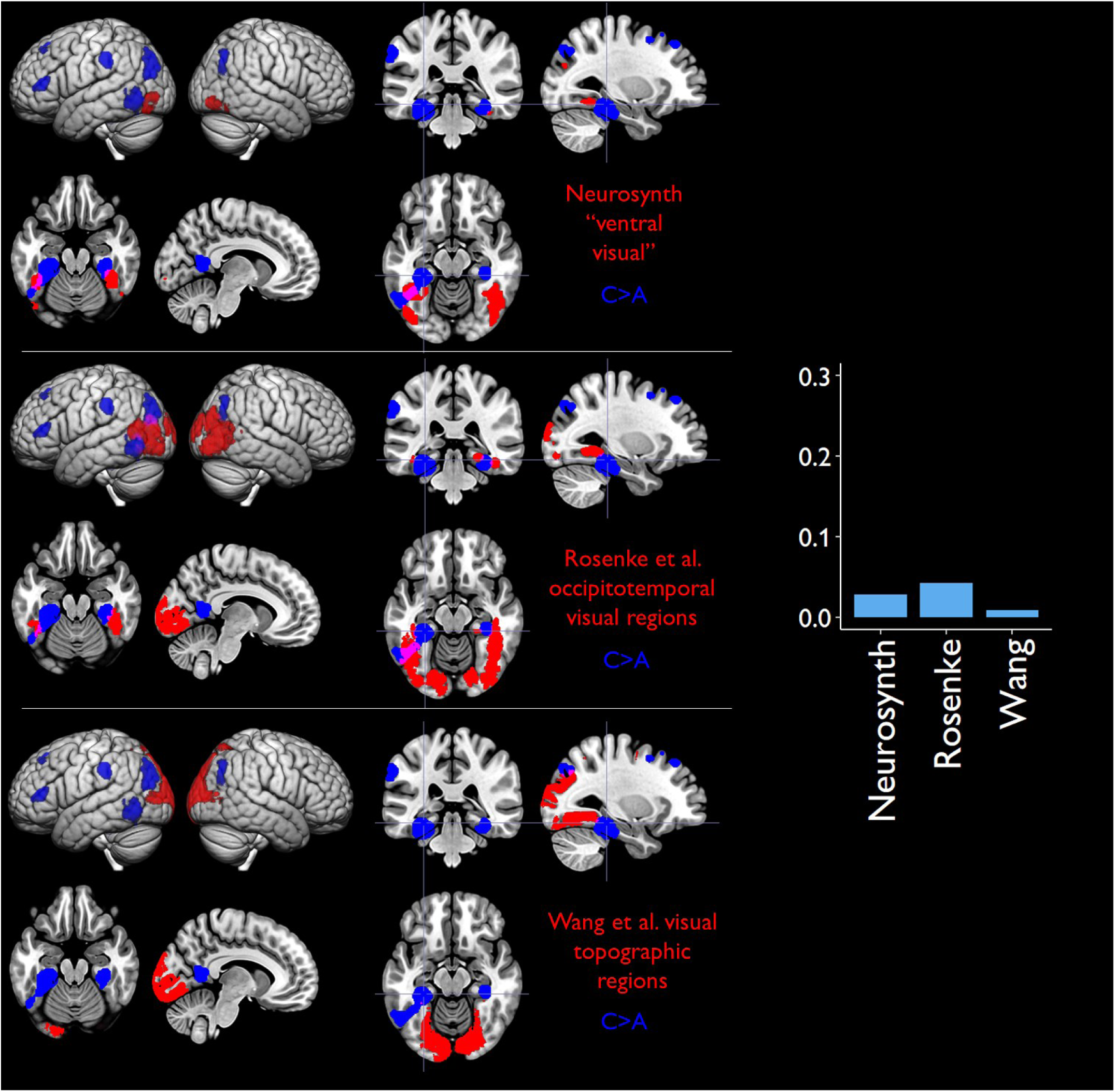
Relationship between C>A effects and alternative definitions of the visual system.

### Supplementary Analysis: Exclusion of episodic memory studies

We repeated the main analysis excluding 7 studies in which participants were instructed to either encode the stimuli into memory or retrieve items from memory during scanning. The results, shown below, were very similar to the main analysis.

**Supplementary Figure 2:**
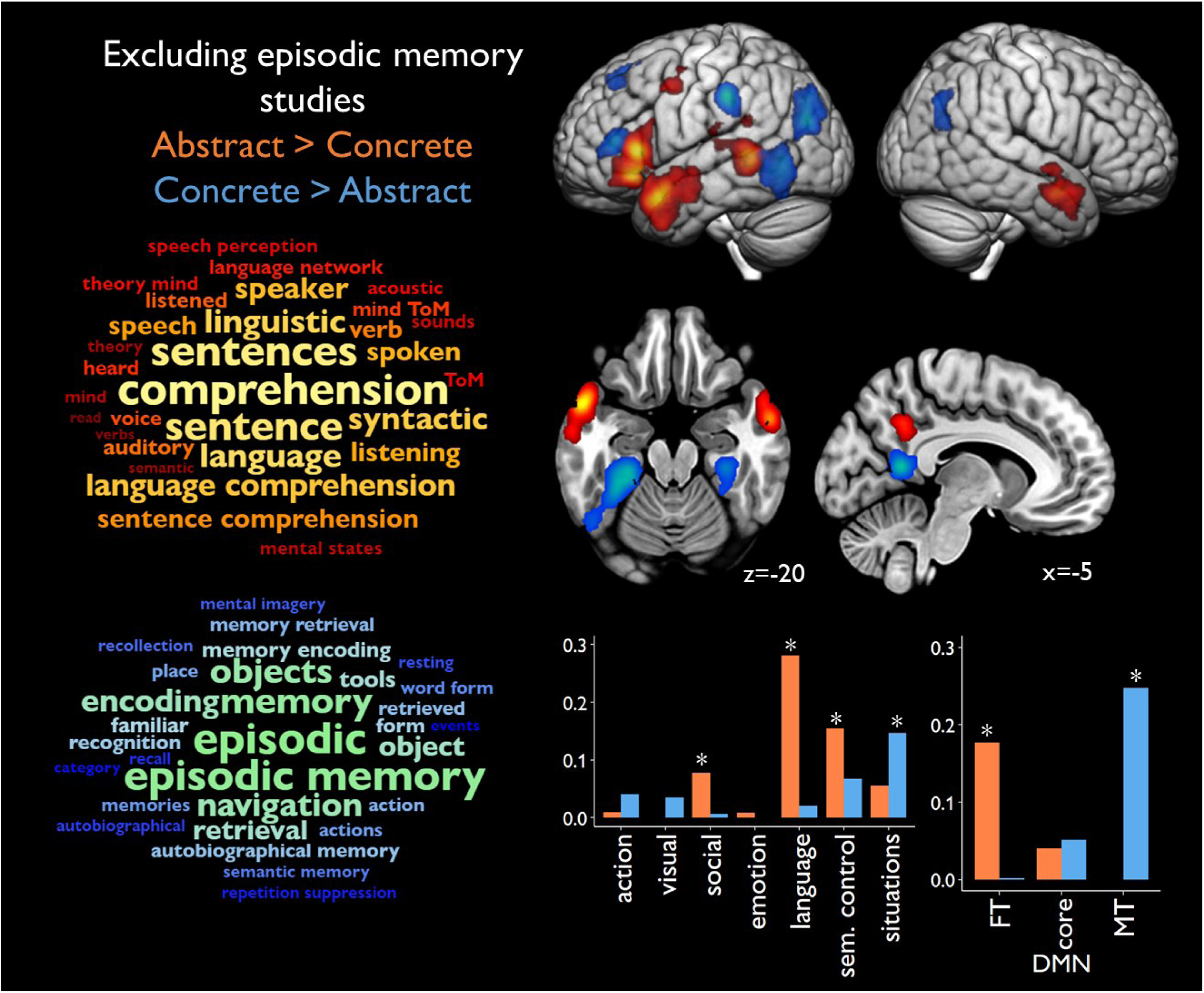
Results of meta-analysis excluding 7 studies that used episodic memory tasks.

## Footnotes

1 Although concreteness is a continuous variable, the vast majority of neuroimaging studies have used factorial manipulations, contrasting a set of highly concrete stimuli with a set of more abstract stimuli. In keeping with this tradition, we use the language of concrete vs. abstract concepts in this paper, while acknowledging that the effects we discuss are likely graded rather than absolute.

## References

Addis, D. R. (2020). Mental Time Travel? A Neurocognitive Model of Event Simulation. Review of Philosophy and Psychology, 11(2), 233–259. 10.1007/s13164-020-00470-0

Alexander-Bloch, A. F., Shou, H., Liu, S., Satterthwaite, T. D., Glahn, D. C., Shinohara, R. T., Vandekar, S. N., & Raznahan, A. (2018). On testing for spatial correspondence between maps of human brain structure and function. NeuroImage, 178, 540–551. 10.1016/j.neuroimage.2018.05.070

Andrews-Hanna, J. R. (2012). The Brain’s Default Network and Its Adaptive Role in Internal Mentation. The Neuroscientist, 18(3), 251–270. 10.1177/1073858411403316

Andrews-Hanna, J. R., & Grilli, M. D. (2021). Mapping the Imaginative Mind: Charting New Paths Forward. Current Directions in Psychological Science, 30(1), 82–89. 10.1177/0963721420980753

Andrews-Hanna, J. R., Reidler, J. S., Sepulcre, J., Poulin, R., & Buckner, R. L. (2010). Functional-anatomic fractionation of the brain’s default network. Neuron, 65(4), 550–562. 10.1016/j.neuron.2010.02.005

Andrews-Hanna, J. R., Smallwood, J., & Spreng, R. N. (2014). The default network and self-generated thought: Component processes, dynamic control, and clinical relevance. Annals of the New York Academy of Sciences, 1316(1), 29. 10.1111/nyas.12360

Auger, S. D., & Maguire, E. A. (2018). Retrosplenial Cortex Indexes Stability beyond the Spatial Domain. The Journal of Neuroscience, 38(6), 1472–1481. 10.1523/JNEUROSCI.2602-17.2017

Baldassano, C., Hasson, U., & Norman, K. A. (2018). Representation of real-world event schemas during narrative perception. Journal of Neuroscience, 38(45), 9689–9699. 10.1523/jneurosci.0251-18.2018

Barsalou, L. W. (1999). Perceptual symbol systems. Behavioral and Brain Sciences, 22(04), 577– 660.

Barsalou, L. W. (2008). Grounded Cognition. Annual Review of Psychology, 59, 617–645. 10.1146/annurev.psych.59.103006.093639

Barsalou, L. W., Dutriaux, L., & Scheepers, C. (2018). Moving beyond the distinction between concrete and abstract concepts. Philosophical Transactions of the Royal Society B: Biological Sciences, 373(1752), 20170144. 10.1098/rstb.2017.0144

Barsalou, L. W., Santos, A., Simmons, W. K., & Wilson, C. D. (2008). Language and simulation in conceptual processing. In A. M. De Vega, A. C. Glenberg, & A. Graesser (Eds.), Symbols, Embodiment and Meaning (pp. 245–283). Oxford University Press.

Barsalou, L. W., & Wiemer-Hastings, K. (2005). Situating Abstract Concepts. In D. Pecher & R. A. Zwaan (Eds.), Grounding Cognition (1st ed., pp. 129–163). Cambridge University Press. 10.1017/CBO9780511499968.007

Bechtold, L., Bellebaum, C., Hoffman, P., & Ghio, M. (2021). Corroborating behavioral evidence for the interplay of representational richness and semantic control in semantic word processing. Scientific Reports, 11(1), Article 1. 10.1038/s41598-021-85711-7

Bedny, M., & Thompson-Schill, S. L. (2006). Neuroanatomically separable effects of imageability and grammatical class during single-word comprehension. Brain and Language, 98(2), 127–139. 10.1016/j.bandl.2006.04.008

Binder, J. R. (2016). In defense of abstract conceptual representations. Psychonomic Bulletin & Review, 23(4), 1096–1108. 10.3758/s13423-015-0909-1

Binder, J. R., Conant, L. L., Humphries, C. J., Fernandino, L., Simons, S. B., Aguilar, M., & Desai, R. H. (2016). Toward a brain-based componential semantic representation. Cognitive Neuropsychology, 33(3–4), 130–174. 10.1080/02643294.2016.1147426

Binder, J. R., & Desai, R. H. (2011). The neurobiology of semantic memory. Trends in Cognitive Sciences, 15(11), 527–536. 10/djtc3r

Binder, J. R., Desai, R. H., Graves, W. W., & Conant, L. L. (2009). Where is the semantic system? A critical review and meta-analysis of 120 functional neuroimaging studies. Cerebral Cortex, *19*, 2767–2796. 10.1093/cercor/bhp055

Binder, J. R., Medler, D. A., Desai, R., Conant, L. L., & Liebenthal, E. (2005). Some neurophysiological constraints on models of word naming. NeuroImage, 27(3), 677–693. 10.1016/j.neuroimage.2005.04.029

Binder, J. R., Westbury, C. F., McKiernan, K. A., Possing, E. T., & Medler, D. A. (2005). Distinct Brain Systems for Processing Concrete and Abstract Concepts. Journal of Cognitive Neuroscience, 17(6), 905–917. 10.1162/0898929054021102

Binney, R. J., Hoffman, P., & Lambon Ralph, M. A. (2016). Mapping the Multiple Graded Contributions of the Anterior Temporal Lobe Representational Hub to Abstract and Social Concepts: Evidence from Distortion-corrected fMRI. Cerebral Cortex, 26(11), 4227–4241. 10.1093/cercor/bhw260

Bird, H., Franklin, S., & Howard, D. (2001). Age of acquisition and imageability ratings for a large set of words, including verbs and function words. Behavior Research Methods Instruments & Computers, 33(1), 73–79. 10/dkxq43

Borghi, A. M., Barca, L., Binkofski, F., Castelfranchi, C., Pezzulo, G., & Tummolini, L. (2019). Words as social tools: Language, sociality and inner grounding in abstract concepts. Physics of Life Reviews, 29, 120–153.

Brysbaert, M., Warriner, A. B., & Kuperman, V. (2014). Concreteness ratings for 40 thousand generally known English word lemmas. Behavior Research Methods, 46(3), 904–911. 10/gd6hzk

Buckner, R. L., Andrews-Hanna, J. R., & Schacter, D. L. (2008). The brain’s default network: Anatomy, function and relevance to disease. Annals of the New York Academy of Sciences, 1124, 1–38. 10.1196/annals.1440.011

Bucur, M., & Papagno, C. (2021). An ALE meta-analytical review of the neural correlates of abstract and concrete words. Scientific Reports, 11, 15727. 10/gs4tfk

Caplan, J. B., & Madan, C. R. (2016). Word Imageability Enhances Association-memory by Increasing Hippocampal Engagement. Journal of Cognitive Neuroscience, 28(10), 1522– 1538. 10.1162/jocn_a_00992

Choudhury, M., Steines, M., Nagels, A., Riedl, L., Kircher, T., & Straube, B. (2021). Neural Basis of Speech-Gesture Mismatch Detection in Schizophrenia Spectrum Disorders. Schizophrenia Bulletin, 47(6), 1761–1771. 10.1093/schbul/sbab059

Chouinard, P. A., & Goodale, M. A. (2010). Category-specific neural processing for naming pictures of animals and naming pictures of tools: An ALE meta-analysis. Neuropsychologia, 48(2), 409–418. 10/fm3drn

Clark, I. A., Kim, M., & Maguire, E. A. (2018). Verbal Paired Associates and the Hippocampus: The Role of Scenes. Journal of Cognitive Neuroscience, 30(12), 1821–1845. 10.1162/jocn_a_01315

Clark, J. M., & Paivio, A. (2004). Extensions of the Paivio, Yuille, and Madigan (1968) norms. Behavior Research Methods Instruments & Computers, *36*(3), 371–383. 10.3758/bf03195584

Coltheart, M. (1981). The MRC Psycholinguistic Database. Quarterly Journal of Experimental Psychology Section A: Human Experimental Psychology, 33(Nov), 497–505. 10.1080/14640748108400805

Conca, F., Borsa, V. M., Cappa, S. F., & Catricalà, E. (2021). The multidimensionality of abstract concepts: A systematic review. Neuroscience & Biobehavioral Reviews, 127, 474–491. 10.1016/j.neubiorev.2021.05.004

Connell, L. (2019). What have labels ever done for us? The linguistic shortcut in conceptual processing. Language, Cognition and Neuroscience, 34(10), 1308–1318. 10.1080/23273798.2018.1471512

Connell, L., & Lynott, D. (2012). Strength of perceptual experience predicts word processing performance better than concreteness or imageability. Cognition, 125(3), 452–465. 10/f4dvqc

Connell, L., Lynott, D., & Banks, B. (2018). Interoception: The forgotten modality in perceptual grounding of abstract and concrete concepts. Philosophical Transactions of the Royal Society B: Biological Sciences, 373(1752), 20170143. 10.1098/rstb.2017.0143

Cortese, M. J., & Fugett, A. (2004). Imageability ratings for 3,000 monosyllabic words. Behavior Research Methods Instruments & Computers, 36(3), 384–387. 10.3758/bf03195585

Cousins, K. A., York, C., Bauer, L., & Grossman, M. (2016). Cognitive and anatomic double dissociation in the representation of concrete and abstract words in semantic variant and behavioral variant frontotemporal degeneration. Neuropsychologia, 84, 244–251. 10/f8m24b

Crutch, S. J., & Warrington, E. K. (2005). Abstract and concrete concepts have structurally different representational frameworks. Brain, 128, 615–627. 10/cqfm9m

D’Argembeau, A., Cassol, H., Phillips, C., Balteau, E., Salmon, E., & Van der Linden, M. (2014). Brains creating stories of selves: The neural basis of autobiographical reasoning. Social Cognitive and Affective Neuroscience, 9(5), 646–652. 10.1093/scan/nst028

Davis, C. P., Altmann, G. T. M., & Yee, E. (2020). Situational systematicity: A role for schema in understanding the differences between abstract and concrete concepts. Cognitive Neuropsychology, 37(1–2), 142–153. 10.1080/02643294.2019.1710124

Della Rosa, P. A., Catricalà, E., Canini, M., Vigliocco, G., & Cappa, S. F. (2018). The left inferior frontal gyrus: A neural crossroads between abstract and concrete knowledge. NeuroImage, 175, 449–459. 10.1016/j.neuroimage.2018.04.021

Della Rosa, P. A., Catricalà, E., Vigliocco, G., & Cappa, S. F. (2010). Beyond the abstract—concrete dichotomy: Mode of acquisition, concreteness, imageability, familiarity, age of acquisition, context availability, and abstractness norms for a set of 417 Italian words. Behavior Research Methods, 42(4), 1042–1048. 10.3758/BRM.42.4.1042

Desai, R. H. (2022). Are metaphors embodied? The neural evidence. Psychological Research, 86(8), 2417–2433. 10/gnw5s5

Desai, R. H., Binder, J. R., Conant, L. L., Mano, Q. R., & Seidenberg, M. S. (2011). The Neural Career of Sensory-motor Metaphors. Journal of Cognitive Neuroscience, 23(9), 2376–2386. 10.1162/jocn.2010.21596

Desai, R. H., Binder, J. R., Conant, L. L., & Seidenberg, M. S. (2010). Activation of Sensory-Motor Areas in Sentence Comprehension. Cerebral Cortex, 20(2), 468–478. 10.1093/cercor/bhp115

Desai, R. H., Conant, L. L., Binder, J. R., Park, H., & Seidenberg, M. S. (2013). A piece of the action: Modulation of sensory-motor regions by action idioms and metaphors. NeuroImage, 83, 862–869. 10.1016/j.neuroimage.2013.07.044

D’Esposito, M., Detre, J. A., Aguirre, G. K., Stallcup, M., Alsop, D. C., Tippet, L. J., & Farah, M. J. (1997). A functional MRI study of mental image generation. Neuropsychologia, 35(5), 725– 730. 10.1016/S0028-3932(96)00121-2

Devlin, J., Chang, M.-W., Lee, K., & Toutanova, K. (2019). *BERT: Pre-training of Deep Bidirectional Transformers for Language Understanding* (arXiv:1810.04805). arXiv. 10.48550/arXiv.1810.04805

Devlin, J. T., Russell, R. P., Davis, M. H., Price, C. J., Moss, H. E., Fadili, M. J., & Tyler, L. K. (2002). Is there an anatomical basis for category-specificity? Semantic memory studies in PET and fMRI. Neuropsychologia, 40(1), 54–75. 10.1016/s0028-3932(01)00066-5

Di Cesare, G., Errante, A., Marchi, M., & Cuccio, V. (2017). Language for action: Motor resonance during the processing of human and robotic voices. Brain and Cognition, 118, 118–127. 10.1016/j.bandc.2017.08.001

Dove, G., Barca, L., Tummolini, L., & Borghi, A. M. (2022). Words have a weight: Language as a source of inner grounding and flexibility in abstract concepts. Psychological Research, 86(8), 2451–2467. 10.1007/s00426-020-01438-6

Eickhoff, S. B., Bzdok, D., Laird, A. R., Kurth, F., & Fox, P. T. (2012). Activation likelihood estimation meta-analysis revisited. Neuroimage, 59(3), 2349–2361. 10.1016/j.neuroimage.2011.09.017

Eickhoff, S. B., Laird, A. R., Grefkes, C., Wang, L. E., Zilles, K., & Fox, P. T. (2009). Coordinate-based activation likelihood estimation meta-analysis of neuroimaging data: A random-effects approach based on empirical estimates of spatial uncertainty. Human Brain Mapping, 30(9), 2907–2926. 10.1002/hbm.20718

Epstein, R. A., & Baker, C. I. (2019). Scene Perception in the Human Brain. Annual Review of Vision Science, 5(Volume 5, 2019), 373–397. 10.1146/annurev-vision-091718-014809

Fernandino, L., & Binder, J. R. (2024). How does the “default mode” network contribute to semantic cognition? Brain and Language, 252, 105405. 10.1016/j.bandl.2024.105405

Fernandino, L., Binder, J. R., Desai, R. H., Pendl, S. L., Humphries, C. J., Gross, W. L., Conant, L. L., & Seidenberg, M. S. (2016). Concept representation reflects multimodal abstraction: A framework for embodied semantics. Cerebral Cortex, 26(5), 2018–2034. 10.1093/cercor/bhv020

Fiebach, C. J., & Friederici, A. D. (2004). Processing concrete words: FMRI evidence against a specific right-hemisphere involvement. Neuropsychologia, 42(1), 62–70. 10.1016/S0028-3932(03)00145-3

Fletcher, P. C., Shallice, T., Frith, C. D., Frackowiak, R. S. J., & Dolan, R. J. (1996). Brain activity during memory retrieval: The influence of imagery and semantic cueing. Brain, 119(5), 1587–1596. 10.1093/brain/119.5.1587

Fliessbach, K., Weis, S., Klaver, P., Elger, C. E., & Weber, B. (2006). The effect of word concreteness on recognition memory. NeuroImage, 32(3), 1413–1421. 10.1016/j.neuroimage.2006.06.007

Gallese, V., & Lakoff, G. (2005). The brain’s concepts: The role of the sensory-motor system in conceptual knowledge. Cognitive Neuropsychology, 22(3–4), 455–479. 10.1080/02643290442000310

Garbarini, F., Calzavarini, F., Diano, M., Biggio, M., Barbero, C., Radicioni, D. P., Geminiani, G., Sacco, K., & Marconi, D. (2020). Imageability effect on the functional brain activity during a naming to definition task. Neuropsychologia, 137, 107275. 10.1016/j.neuropsychologia.2019.107275

Ghio, M., Vaghi, M. M. S., Perani, D., & Tettamanti, M. (2016). Decoding the neural representation of fine-grained conceptual categories. NeuroImage, 132, 93–103. 10.1016/j.neuroimage.2016.02.009

Giacobbe, C., Raimo, S., Cropano, M., & Santangelo, G. (2022). Neural correlates of embodied action language processing: A systematic review and meta-analytic study. Brain Imaging and Behavior, 16(5), 2353–2374. 10.1007/s11682-022-00680-3

Giallanza, T., Campbell, D., Cohen, J. D., & Rogers, T. (2023). An Integrated Model of Semantics and Control. PsyArXiv. 10.31234/osf.io/jq7ta

Gibbs, R. W. (1994). The poetics of mind: Figurative thought, language, and understanding. Cambridge University Press.

Giesbrecht, B. (2004). Separable Effects of Semantic Priming and Imageability on Word Processing in Human Cortex. Cerebral Cortex, 14(5), 521–529. 10.1093/cercor/bhh014

Gilboa, A., & Marlatte, H. (2017). Neurobiology of Schemas and Schema-Mediated Memory. Trends in Cognitive Sciences, 21(8), 618–631. 10.1016/j.tics.2017.04.013

Gilead, M., Liberman, N., & Maril, A. (2013). The language of future-thought: An fMRI study of embodiment and tense processing. NeuroImage, 65, 267–279. 10.1016/j.neuroimage.2012.09.073

Glenberg, A. M., & Robertson, D. A. (2000). Symbol grounding and meaning: A comparison of high-dimensional and embodied theories of meaning. Journal of Memory and Language, 43(3), 379–401. 10/dsr5gr

Goldberg, R. F., Perfetti, C. A., Fiez, J. A., & Schneider, W. (2007). Selective Retrieval of Abstract Semantic Knowledge in Left Prefrontal Cortex. The Journal of Neuroscience, 27(14), 3790– 3798. 10.1523/JNEUROSCI.2381-06.2007

Graves, W. W., Desai, R., Humphries, C., Seidenberg, M. S., & Binder, J. R. (2010). Neural Systems for Reading Aloud: A Multiparametric Approach. Cerebral Cortex, 20(8), 1799–1815. 10.1093/cercor/bhp245

Grossman, M., Koenig, P., DeVita, C., Glosser, G., Alsop, D., Detre, J., & Gee, J. (2002). The Neural Basis for Category-Specific Knowledge: An fMRI Study. NeuroImage, 15(4), 936– 948. 10.1006/nimg.2001.1028

Gurguryan, L., & Sheldon, S. (2019). Retrieval orientation alters neural activity during autobiographical memory recollection. NeuroImage, 199, 534–544. 10.1016/j.neuroimage.2019.05.077

Harris, G. J., Chabris, C. F., Clark, J., Urban, T., Aharon, I., Steele, S., McGrath, L., Condouris, K., & Tager-Flusberg, H. (2006). Brain activation during semantic processing in autism spectrum disorders via functional magnetic resonance imaging. Brain and Cognition, 61(1), 54–68. 10.1016/j.bandc.2005.12.015

Hauk, O., Davis, M. H., Kherif, F., & Pulvermüller, F. (2008). Imagery or meaning? Evidence for a semantic origin of category-specific brain activity in metabolic imaging. European Journal of Neuroscience, 27(7), 1856–1866. 10.1111/j.1460-9568.2008.06143.x

Hauk, O., Johnsrude, I., & Pulvermuller, F. (2004). Somatotopic representation of action words in human motor and premotor cortex. Neuron, 41(2), 301–307. 10.1016/s0896-6273(03)00838-9

Hayashi, A., Okamoto, Y., Yoshimura, S., Yoshino, A., Toki, S., Yamashita, H., Matsuda, F., & Yamawaki, S. (2014). Visual imagery while reading concrete and abstract Japanese kanji words: An fMRI study. Neuroscience Research, 79, 61–66. 10.1016/j.neures.2013.10.007

Heidlmayr, K., Weber, K., Takashima, A., & Hagoort, P. (2020). No title, no theme: The joined neural space between speakers and listeners during production and comprehension of multi-sentence discourse. Cortex, *130*, 111–126. 10.1016/j.cortex.2020.04.035

Hoffman, P. (2016). The meaning of ‘life’ and other abstract words: Insights from neuropsychology. J Neuropsychol, 10, 317–343. 10.1111/jnp.12065

Hoffman, P., Binney, R. J., & Lambon Ralph, M. A. (2015). Differing contributions of inferior prefrontal and anterior temporal cortex to concrete and abstract conceptual knowledge. Cortex, 63, 250–265. 10.1016/j.cortex.2014.09.001

Hoffman, P., Jefferies, E., & Lambon Ralph, M. A. (2010). Ventrolateral prefrontal cortex plays an executive regulation role in comprehension of abstract words: Convergent neuropsychological and repetitive TMS evidence. Journal of Neuroscience, 30(46), 15450– 15456. 10.1523/jneurosci.3783-10.2010

Hoffman, P., & Lambon Ralph, M. A. (2013). Shapes, scents and sounds: Quantifying the full multi- sensory basis of conceptual knowledge. Neuropsychologia, 51(1), 14–25. 10/f4mns4

Hoffman, P., Lambon Ralph, M. A., & Rogers, T. T. (2013). Semantic diversity: A measure of semantic ambiguity based on variability in the contextual usage of words. Behavior Research Methods, 45(3), 718–730. 10.3758/s13428-012-0278-x

Hoffman, P., & Morcom, A. M. (2018). Age-related changes in the neural networks supporting semantic cognition: A meta-analysis of 47 functional neuroimaging studies. Neuroscience & Biobehavioral Reviews, 84, 134–150. 10/gctxfv

Jackson, R. L. (2021). The neural correlates of semantic control revisited. Neuroimage, 224, 117444. 10.1016/j.neuroimage.2020.117444

Jessen, F., Heun, R., Erb, M., Granath, D.-O., Klose, U., Papassotiropoulos, A., & Grodd, W. (2000). The Concreteness Effect: Evidence for Dual Coding and Context Availability. Brain and Language, 74(1), 103–112. 10.1006/brln.2000.2340

Johnson-Laird, P. N. (1988). The computer and the mind: An introduction to cognitive science. Harvard University Press.

Joue, G., Boven, L., Willmes, K., Evola, V., Demenescu, L. R., Hassemer, J., Mittelberg, I., Mathiak, K., Schneider, F., & Habel, U. (2020). Metaphor processing is supramodal semantic processing: The role of the bilateral lateral temporal regions in multimodal communication. Brain and Language, 205, 104772. 10.1016/j.bandl.2020.104772

Kana, R. K., Blum, E. R., Ladden, S. L., & Ver Hoef, L. W. (2012). “How to do things with Words”: Role of motor cortex in semantic representation of action words. Neuropsychologia, 50(14), 3403–3409. 10.1016/j.neuropsychologia.2012.09.006

Kiehl, K. A., Liddle, P. F., Smith, A. M., Mendrek, A., Forster And, B. B., & Hare, R. D. (1999). Neural pathways involved in the processing of concrete and abstract words. Human Brain Mapping, 7(4), 225–233. 10.1002/(SICI)1097-0193(1999)7:4<225::AID-HBM1>3.0.CO;2-P

Kounios, J., Koenig, P., Glosser, G., DeVita, C., Dennis, K., Moore, P., & Grossman, M. (2003). Category-specific medial temporal lobe activation and the consolidation of semantic memory: Evidence from fMRI. Cognitive Brain Research, 17(2), 484–494. 10.1016/S0926-6410(03)00164-2

Kousta, S. T., Vigliocco, G., Vinson, D. P., Andrews, M., & Del Campo, E. (2011). The Representation of Abstract Words: Why Emotion Matters. J Exp Psychol Gen, 140(1), 14–34. 10/brjczk

Kumar, U. (2016). Neural dichotomy of word concreteness: A view from functional neuroimaging. Cognitive Processing, 17(1), 39–48. 10.1007/s10339-015-0738-1

Kuperberg, G. R., West, W. C., Lakshmanan, B. M., & Goff, D. (2008). Functional Magnetic Resonance Imaging Reveals Neuroanatomical Dissociations During Semantic Integration in Schizophrenia. Biological Psychiatry, 64(5), 407–418. 10.1016/j.biopsych.2008.03.018

Lakoff, G., & Johnson, M. (1999). Philosophy in the flesh: The embodied mind and its challenge to western thought. Basic Books.

Lambon Ralph, M. A., Jefferies, E., Patterson, K., & Rogers, T. T. (2017). The neural and computational bases of semantic cognition. Nature Reviews Neuroscience, 18, 42–55. 10/gmhhtx

Lee, S., Parthasarathi, T., & Kable, J. W. (2021). The Ventral and Dorsal Default Mode Networks Are Dissociably Modulated by the Vividness and Valence of Imagined Events. Journal of Neuroscience, 41(24), 5243–5250. 10.1523/JNEUROSCI.1273-20.2021

Lenci, A., Lebani, G. E., & Passaro, L. C. (2018). The emotions of abstract words: A distributional semantic analysis. Topics in Cognitive Science, 10(3), 550–572. 10.1111/tops.12335

Lin, N., Wang, X., Xu, Y., Wang, X., Hua, H., Zhao, Y., & Li, X. (2018). Fine Subdivisions of the Semantic Network Supporting Social and Sensory–Motor Semantic Processing. Cerebral Cortex, 28(8), 2699–2710. 10.1093/cercor/bhx148

Louwerse, M. M. (2011). Symbol interdependency in symbolic and embodied cognition. Topics in Cognitive Science, 3(2), 273–302. 10.1111/j.1756-8765.2010.01106.x

Lynott, D., Connell, L., Brysbaert, M., Brand, J., & Carney, J. (2020). The Lancaster Sensorimotor Norms: Multidimensional measures of perceptual and action strength for 40,000 English words. Behavior Research Methods, 52(3), 1271–1291. 10.3758/s13428-019-01316-z

Margulies, D. S., Ghosh, S. S., Goulas, A., Falkiewicz, M., Huntenburg, J. M., Langs, G., Bezgin, G., Eickhoff, S. B., Castellanos, F. X., & Petrides, M. (2016). Situating the default-mode network along a principal gradient of macroscale cortical organization. Proceedings of the National Academy of Sciences, 113(44), 12574–12579. 10.1073/pnas.1608282113

Markello, R. D., Hansen, J. Y., Liu, Z.-Q., Bazinet, V., Shafiei, G., Suárez, L. E., Blostein, N., Seidlitz, J., Baillet, S., Satterthwaite, T. D., Chakravarty, M. M., Raznahan, A., & Misic, B. (2022). neuromaps: Structural and functional interpretation of brain maps. Nature Methods, 19(11), 1472–1479. 10.1038/s41592-022-01625-w

Martin, A. (2016). GRAPES—Grounding representations in action, perception, and emotion systems: How object properties and categories are represented in the human brain. Psychonomic Bulletin & Review, 23, 979–990. 10.3758/s13423-015-0842-3

Martin, A., Haxby, J. V., Lalonde, F. M., Wiggs, C. L., & Ungerleider, L. G. (1995). Discrete cortical regions associated with knowledge of color and knowledge of action. Science, 270, 102–105. 10/fv2rf4

Mayer, K. M., Macedonia, M., & Von Kriegstein, K. (2017). Recently learned foreign abstract and concrete nouns are represented in distinct cortical networks similar to the native language: Abstract and Concrete Nouns in the Brain. Human Brain Mapping, 38(9), 4398–4412. 10.1002/hbm.23668

Mckiernan, K. A., Kaufman, J. N., Kucera-Thompson, J., & Binder, J. R. (2003). A parametric manipulation of factors affecting task-induced deactivation in functional neuroimaging. Journal of Cognitive Neuroscience, 15(3), 394–408. 10/b6j6ht

McRae, K., Nedjadrasul, D., Pau, R., Lo, B. P.-H., & King, L. (2018). Abstract Concepts and Pictures of Real-World Situations Activate One Another. Topics in Cognitive Science, 10(3), 518–532. 10.1111/tops.12328

Meersmans, K., Bruffaerts, R., Jamoulle, T., Liuzzi, A. G., De Deyne, S., Storms, G., Dupont, P., & Vandenberghe, R. (2020). Representation of associative and affective semantic similarity of abstract words in the lateral temporal perisylvian language regions. NeuroImage, 217, 116892. 10.1016/j.neuroimage.2020.116892

Meersmans, K., Storms, G., De Deyne, S., Bruffaerts, R., Dupont, P., & Vandenberghe, R. (2022). Orienting to different dimensions of word meaning alters the representation of word meaning in early processing regions. Cerebral Cortex, 32(15), 3302–3317. 10.1093/cercor/bhab416

Mellet, E., Tzourio, N., Denis, M., & Mazoyer, B. (1998). Cortical anatomy of mental imagery of concrete nouns based on their dictionary definition: NeuroReport, 9(5), 803–808. 10.1097/00001756-199803300-00007

Mestres-Missé, A., Münte, T. F., & Rodriguez-Fornells, A. (2009). Functional Neuroanatomy of Contextual Acquisition of Concrete and Abstract Words. Journal of Cognitive Neuroscience, 21(11), 2154–2171. 10.1162/jocn.2008.21171

Mikolov, T., Chen, K., Corrado, G., & Dean, J. (2013). Efficient estimation of word representations in vector space. ArXiv Preprint ArXiv:1301.3781.

Morales, M., Patel, T., Tamm, A., Pickering, M., & Hoffman, P. (2022). Similar neural networks respond to coherence during comprehension and production of discourse. Cerebral Cortex, 32(19), 4317–4330. 10.1093/cercor/bhab485

Mullally, S. L., & Maguire, E. A. (2014). Memory, Imagination, and Predicting the Future: A Common Brain Mechanism? The Neuroscientist, 20(3), 220–234. 10.1177/1073858413495091

Nagels, A., Chatterjee, A., Kircher, T., & Straube, B. (2013). The role of semantic abstractness and perceptual category in processing speech accompanied by gestures. Frontiers in Behavioral Neuroscience, 7. 10.3389/fnbeh.2013.00181

Newcombe, P. I., Campbell, C., Siakaluk, P. D., & Pexman, P. M. (2012). Effects of emotional and sensorimotor knowledge in semantic processing of concrete and abstract nouns. Frontiers in Human Neuroscience, 6, 275. 10.3389/fnhum.2012.00275

Newell, A. (1980). Physical symbol systems. Cognitive Science, 4(2), 135–183.

Noppeney, U., & Price, C. J. (2002). Retrieval of Visual, Auditory, and Abstract Semantics. NeuroImage, 15(4), 917–926. 10.1006/nimg.2001.1016

Noppeney, U., & Price, C. J. (2004). Retrieval of abstract semantics. NeuroImage, 22(1), 164–170. 10.1016/j.neuroimage.2003.12.010

Olivetti Belardinelli, M., Palmiero, M., Sestieri, C., Nardo, D., Di Matteo, R., Londei, A., D’Ausilio, A., Ferretti, A., Del Gratta, C., & Romani, G. L. (2009). An fMRI investigation on image generation in different sensory modalities: The influence of vividness. Acta Psychologica, 132(2), 190–200. 10.1016/j.actpsy.2009.06.009

Paivio, A. (1986). Mental representations: A dual-coding approach. Oxford University Press.

Papeo, L., Rumiati, R. I., Cecchetto, C., & Tomasino, B. (2012). On-line Changing of Thinking about Words: The Effect of Cognitive Context on Neural Responses to Verb Reading. Journal of Cognitive Neuroscience, 24(12), 2348–2362. 10.1162/jocn_a_00291

Patel, T., Morales, M., Pickering, M. J., & Hoffman, P. (2023). A common neural code for meaning in discourse production and comprehension. NeuroImage, 279, 120295. 10.1016/j.neuroimage.2023.120295

Pauligk, S., Kotz, S. A., & Kanske, P. (2019). Differential Impact of Emotion on Semantic Processing of Abstract and Concrete Words: ERP and fMRI Evidence. Scientific Reports, 9(1), 14439. 10.1038/s41598-019-50755-3

Pecher, D. (2018). Curb your embodiment. Topics in Cognitive Science, 10(3), 501–517. 10/gd5pgp

Perani, D., Cappa, S. F., Schnur, T., Tettamanti, M., Collina, S., Rosa, M. M., & Fazio1, F. (1999). The neural correlates of verb and noun processing. Brain, *122*(12), 2337–2344. 10.1093/brain/122.12.2337

Pereira, F., Gershman, S., Ritter, S., & Botvinick, M. (2016). A comparative evaluation of off-the-shelf distributed semantic representations for modelling behavioural data. Cognitive Neuropsychology, 33(3–4), 175–190. 10/ghcv8k

Pexman, P. M., Diveica, V., & Binney, R. J. (2022). Social semantics: The organization and grounding of abstract concepts. Philosophical Transactions of the Royal Society B: Biological Sciences, 378(1870), 20210363. 10.1098/rstb.2021.0363

Pexman, P. M., Hargreaves, I. S., Edwards, J. D., Henry, L. C., & Goodyear, B. G. (2007). Neural Correlates of Concreteness in Semantic Categorization. Journal of Cognitive Neuroscience, 19(8), 1407–1419. 10.1162/jocn.2007.19.8.1407

Poerio, G. L., Sormaz, M., Wang, H.-T., Margulies, D., Jefferies, E., & Smallwood, J. (2017). The role of the default mode network in component processes underlying the wandering mind. Social Cognitive and Affective Neuroscience, 12(7), 1047–1062. 10.1093/scan/nsx041

Pulvermüller, F. (2013). How neurons make meaning: Brain mechanisms for embodied and abstract-symbolic semantics. Trends in Cognitive Sciences, 17(9), 458–470. 10.1016/j.tics.2013.06.004

Quinlan, P. T. (1992). Oxford Psycholinguistic Database. Oxford University Press.

Raichle, M. E., MacLeod, A. M., Snyder, A. Z., Powers, W. J., Gusnard, D. A., & Shulman, G. L. (2001). A default mode of brain function. Proceedings of the National Academy of Sciences, 98(2), 676–682. 10.1073/pnas.98.2.676

Ranganath, C., & Ritchey, M. (2012). Two cortical systems for memory-guided behaviour. Nature Reviews Neuroscience, 13(10), 713–726. 10.1038/nrn3338

Reilly, J., Shain, C., Borghesani, V., Kuhnke, P., Vigliocco, G., Peelle, J., Mahon, B., Buxbaum, L., Majid, A., Brysbaert, M., Borghi, A., De Deyne, S., Dove, G., Papeo, L., Pexman, P., Poeppel, D., Lupyan, G., Boggio, P., Hickock, G., … Vinson, D. (2025). What we mean when we say semantic: Toward a multidisciplinary semantic glossary. Psychonomic Bulletin & Review, 32, 243–280. 10.3758/s13423-024-02556-7

Rodríguez-Ferreiro, J., Gennari, S. P., Davies, R., & Cuetos, F. (2011). Neural Correlates of Abstract Verb Processing. Journal of Cognitive Neuroscience, 23(1), 106–118. 10.1162/jocn.2010.21414

Rosenke, M., van Hoof, R., van den Hurk, J., Grill-Spector, K., & Goebel, R. (2021). A Probabilistic Functional Atlas of Human Occipito-Temporal Visual Cortex. Cerebral Cortex, 31(1), 603– 619. 10.1093/cercor/bhaa246

Roxbury, T., McMahon, K., & Copland, D. A. (2014). An fMRI study of concreteness effects in spoken word recognition. Behavioral and Brain Functions, 10(1), 34. 10.1186/1744-9081-10-34

Rubin, T. N., Koyejo, O., Gorgolewski, K. J., Jones, M. N., Poldrack, R. A., & Yarkoni, T. (2017). Decoding brain activity using a large-scale probabilistic functional-anatomical atlas of human cognition. PLoS Computational Biology, 13(10), e1005649. 10/gcgdtm

Rüschemeyer, S.-A., Brass, M., & Friederici, A. D. (2007). Comprehending Prehending: Neural Correlates of Processing Verbs with Motor Stems. Journal of Cognitive Neuroscience, 19(5), 855–865. 10.1162/jocn.2007.19.5.855

Sabsevitz, D. S., Medler, D. A., Seidenberg, M., & Binder, J. R. (2005). Modulation of the semantic system by word imageability. NeuroImage, 27(1), 188–200. 10.1016/j.neuroimage.2005.04.012

Sakreida, K., Scorolli, C., Menz, M. M., Heim, S., Borghi, A. M., & Binkofski, F. (2013). Are abstract action words embodied? An fMRI investigation at the interface between language and motor cognition. Frontiers in Human Neuroscience, 7. 10.3389/fnhum.2013.00125

Saygin, A. P., McCullough, S., Alac, M., & Emmorey, K. (2010). Modulation of BOLD response in motion-sensitive lateral temporal cortex by real and fictive motion sentences. Journal of Cognitive Neuroscience, 22(11), 2480–2490. 10.1162/jocn.2009.21388

Schwanenflugel, P. J., Harnishfeger, K. K., & Stowe, R. W. (1988). Context Availability and Lexical Decisions for Abstract and Concrete Words. Journal of Memory and Language, 27(5), 499–520. 10/dt3z52

Schwanenflugel, P. J., & Shoben, E. J. (1983). Differential Context Effects in the Comprehension of Abstract and Concrete Verbal Materials. *Journal of Experimental Psychology: Learning*, Memory and Cognition, 9(1), 82–102. 10/d4jwbt

Shafto, M., Randall, B., Stamatakis, E. A., Wright, P., & Tyler, L. K. (2012). Age-related Neural Reorganization during Spoken Word Recognition: The Interaction of Form and Meaning. Journal of Cognitive Neuroscience, 24(6), 1434–1446. 10.1162/jocn_a_00218

Shao, X., Krieger-Redwood, K., Zhang, M., Hoffman, P., Lanzoni, L., Leech, R., Smallwood, J., & Jefferies, E. (2024). Distinctive and complementary roles of default mode network subsystems in semantic cognition. Journal of Neuroscience, 44(20), e1907232024. 10.1523/JNEUROSCI.1907-23.2024

Silson, E. H., Steel, A., Kidder, A., Gilmore, A. W., & Baker, C. I. (2019). Distinct subdivisions of human medial parietal cortex support recollection of people and places. ELife, 8, e47391. 10.7554/eLife.47391

Smallwood, J., Bernhardt, B. C., Leech, R., Bzdok, D., Jefferies, E., & Margulies, D. S. (2021). The default mode network in cognition: A topographical perspective. Nature Reviews Neuroscience, 22(8), 503–513. 10.1038/s41583-021-00474-4

Smallwood, J., Tipper, C., Brown, K., Baird, B., Engen, H., Michaels, J. R., Grafton, S., & Schooler, J. W. (2013). Escaping the here and now: Evidence for a role of the default mode network in perceptually decoupled thought. NeuroImage, 69, 120–125. 10.1016/j.neuroimage.2012.12.012

Stadthagen-Gonzalez, H., & Davis, C. J. (2006). The Bristol norms for age of acquisition, imageability, and familiarity. Behavior Research Methods, 38(4), 598–605. 10.3758/bf03193891

Straube, B., Green, A., Bromberger, B., & Kircher, T. (2011). The differentiation of iconic and metaphoric gestures: Common and unique integration processes. Human Brain Mapping, 32(4), 520–533. 10.1002/hbm.21041

Straube, B., He, Y., Steines, M., Gebhardt, H., Kircher, T., Sammer, G., & Nagels, A. (2013). Supramodal neural processing of abstract information conveyed by speech and gesture. Frontiers in Behavioral Neuroscience, 7. 10.3389/fnbeh.2013.00120

Tamir, D. I., Bricker, A. B., Dodell-Feder, D., & Mitchell, J. P. (2016). Reading fiction and reading minds: The role of simulation in the default network. Social Cognitive and Affective Neuroscience, 11(2), 215–224. 10.1093/scan/nsv114

Tettamanti, M., Buccino, G., Saccuman, M. C., Gallese, V., Danna, M., Scifo, P., Fazio, F., Rizzolatti, G., Cappa, S. F., & Perani, D. (2005). Listening to Action-related Sentences Activates Fronto-parietal Motor Circuits. Journal of Cognitive Neuroscience, 17(2), 273–281. 10.1162/0898929053124965

Tettamanti, M., Manenti, R., Della Rosa, P. A., Falini, A., Perani, D., Cappa, S. F., & Moro, A. (2008). Negation in the brain: Modulating action representations. NeuroImage, 43(2), 358–367. 10.1016/j.neuroimage.2008.08.004

Thompson-Schill, S. L., Aguirre, G. K., D’Esposito, M., & Farah, M. J. (1999). A neural basis for category and modality specificity of semantic knowledge. Neuropsychologia, 37, 671–676. 10/fts7nx

Tian, L., Chen, H., Zhao, W., Wu, J., Zhang, Q., De, A., Leppänen, P., Cong, F., & Parviainen, T. (2020). The role of motor system in action-related language comprehension in L1 and L2: An fMRI study. Brain and Language, 201, 104714. 10.1016/j.bandl.2019.104714

Toglia, M. P., & Battig, W. F. (1978). Handbook of semantic word norms (pp. vii, 152). Lawrence Erlbaum.

Tomasino, B., Fabbro, F., & Brambilla, P. (2014). How do conceptual representations interact with processing demands: An fMRI study on action- and abstract-related words. Brain Research, 1591, 38–52. 10.1016/j.brainres.2014.10.008

Tong, J., Binder, J. R., Humphries, C., Mazurchuk, S., Conant, L. L., & Fernandino, L. (2022). A Distributed Network for Multimodal Experiential Representation of Concepts. The Journal of Neuroscience, 42(37), 7121–7130. 10.1523/JNEUROSCI.1243-21.2022

Troche, J., Crutch, S., & Reilly, J. (2014). Clustering, hierarchical organization, and the topography of abstract and concrete nouns. Frontiers in Psychology, 5, 360. 10/gbf4r5

Turkeltaub, P. E., Eickhoff, S. B., Laird, A. R., Fox, M., Wiener, M., & Fox, P. (2012). Minimizing within-experiment and within-group effects in activation likelihood estimation meta-analyses. Human Brain Mapping, 33(1), 1–13. 10.1002/hbm.21186

Tyler, L. K. (2001). The neural representation of nouns and verbs: PET studies. Brain, 124(8), 1619– 1634. 10.1093/brain/124.8.1619

Van Dam, W. O., & Desai, R. H. (2016). The Semantics of Syntax: The Grounding of Transitive and Intransitive Constructions. Journal of Cognitive Neuroscience, 28(5), 693–709. 10.1162/jocn_a_00926

Van Dam, W. O., Rueschemeyer, S.-A., & Bekkering, H. (2010). How specifically are action verbs represented in the neural motor system: An fMRI study. NeuroImage, 53(4), 1318–1325. 10.1016/j.neuroimage.2010.06.071

van Dam, W. O., van Dijk, M., Bekkering, H., & Rueschemeyer, S. A. (2012). Flexibility in embodied lexical-semantic representations. Human Brain Mapping, 33(10), 2322–2333. 10/bw8sdp

Vigliocco, G., Kousta, S. T., Della Rosa, P. A., Vinson, D. P., Tettamanti, M., Devlin, J. T., & Cappa, S. F. (2014). The neural representation of abstract words: The role of emotion. Cerebral Cortex, 24, 1767–1777. 10.1093/cercor/bht025

Vigliocco, G., Meteyard, L., Andrews, M., & Kousta, S. (2009). Toward a theory of semantic representation. Language and Cognition, 1(02), 219–247. 10/fvwk3v

Villani, C., Loia, A., & Bolognesi, M. M. (2024). The semantic content of concrete, abstract, specific, and generic concepts. Language and Cognition, 16(4), 867–894. 10.1017/langcog.2023.64

Villani, C., Lugli, L., Liuzza, M. T., & Borghi, A. M. (2019). Varieties of abstract concepts and their multiple dimensions. Language and Cognition, 11(3), 403–430. 10.1017/langcog.2019.23

Vitello, S., & Rodd, J. M. (2015). Resolving semantic ambiguities in sentences: Cognitive processes and brain mechanisms. Language and Linguistics Compass, 9(10), 391–405. 10.1111/lnc3.12160

Wallentin, M., Ostergaard, S., Lund, T., Ostergaard, L., & Roepstorff, A. (2005). Concrete spatial language: See what I mean? Brain and Language, 92(3), 221–233. 10.1016/j.bandl.2004.06.106

Wang, J., Conder, J. A., Blitzer, D. N., & Shinkareva, S. V. (2010). Neural Representation of Abstract and Concrete Concepts: A Meta-Analysis of Neuroimaging Studies. Hum Brain Mapp, 31(10), 1459–1468. 10/cw55dk

Wang, L., Mruczek, R. E. B., Arcaro, M. J., & Kastner, S. (2015). Probabilistic Maps of Visual Topography in Human Cortex. Cerebral Cortex, 25(10), 3911–3931. 10.1093/cercor/bhu277

Wang, X., & Bi, Y. (2021). Idiosyncratic tower of Babel: Individual differences in word-meaning representation increase as word abstractness increases. Psychological Science, 32(10), 1617– 1635. 10.1177/09567976211003877

Wang, X., Wang, B., & Bi, Y. (2019). Close yet independent: Dissociation of social from valence and abstract semantic dimensions in the left anterior temporal lobe. Human Brain Mapping, 40(16), 4759–4776. 10.1002/hbm.24735

Wang, X., Wu, W., Ling, Z., Xu, Y., Fang, Y., Wang, X., Binder, J. R., Men, W., Gao, J.-H., & Bi, Y. (2018). Organizational principles of abstract words in the human brain. Cerebral Cortex, 28(12), 4305–4318. 10.1093/cercor/bhx283

Watson, C. E., Cardillo, E. R., Ianni, G. R., & Chatterjee, A. (2013). Action concepts in the brain: An activation likelihood estimation meta-analysis. Journal of Cognitive Neuroscience, 25(8), 1191–1205. 10/ghms3p

Westbury, C. F., Cribben, I., & Cummine, J. (2016). Imaging Imageability: Behavioral Effects and Neural Correlates of Its Interaction with Affect and Context. Frontiers in Human Neuroscience, 10. 10.3389/fnhum.2016.00346

Whatmough, C., Verret, L., Fung, D., & Chertkow, H. (2004). Common and Contrasting Areas of Activation for Abstract and Concrete Concepts: An H215O PET Study. Journal of Cognitive Neuroscience, 16(7), 1211–1226. 10.1162/0898929041920540

Wiemer-Hastings, K., & Xu, X. (2005). Content differences for abstract and concrete concepts. Cognitive Science, 29(5), 719–736. 10/cd4zxg

Winter, B. (2023). Abstract concepts and emotion: Cross-linguistic evidence and arguments against affective embodiment. Philosophical Transactions of the Royal Society B, 378(1870), 20210368. 10/gs4td3

Wise, R. J. S., Howard, D., Mummery, C. J., Fletcher, P., Le, A., & Scott, S. K. (2000). Noun imageability and the temporal lobes. Neuropsychologia, 38(7), 985–994. 10.1016/S0028-3932(99)00152-9

Yarkoni, T., Poldrack, R. A., Nichols, T. E., Van Essen, D. C., & Wager, T. D. (2011). Large-scale automated synthesis of human functional neuroimaging data. Nature Methods, 8(8), 665–670. 10.1038/nmeth.1635

Yeo, B. T. T., Krienen, F. M., Sepulcre, J., Sabuncu, M. R., Lashkari, D., Hollinshead, M., Roffman, J. L., Smoller, J. W., Zöllei, L., Polimeni, J. R., Fischl, B., Liu, H., & Buckner, R. L. (2011). The organization of the human cerebral cortex estimated by intrinsic functional connectivity. Journal of Neurophysiology, 106(3), 1125–1165. 10.1152/jn.00338.2011

Yeshurun, Y., Nguyen, M., & Hasson, U. (2021). The default mode network: Where the idiosyncratic self meets the shared social world. Nature Reviews Neuroscience, 22(3), 181–192. 10.1038/s41583-020-00420-w

Yu, S., Gu, C., Huang, K., & Li, P. (2024). Predicting the next sentence (not word) in large language models: What model-brain alignment tells us about discourse comprehension. Science Advances, 10(21), eadn7744. 10.1126/sciadv.adn7744

Zeki, S. (2004). Thirty years of a very special visual area, Area V5. The Journal of Physiology, *557*(Pt 1), 1–2. 10.1113/jphysiol.2004.063040

Zhuang, J., Randall, B., Stamatakis, E. A., Marslen-Wilson, W. D., & Tyler, L. K. (2011). The Interaction of Lexical Semantics and Cohort Competition in Spoken Word Recognition: An fMRI Study. Journal of Cognitive Neuroscience, 23(12), 3778–3790. 10.1162/jocn_a_00046

Zwaan, R. A. (2014). Embodiment and language comprehension: Reframing the discussion. Trends in Cognitive Sciences, 18(5), 229–234. 10.1016/j.tics.2014.02.008

